# Efficient inverse graphics in biological face processing

**DOI:** 10.1101/282798

**Authors:** Ilker Yildirim, Mario Belledonne, Winrich Freiwald, Joshua Tenenbaum

**Affiliations:** Department of Brain & Cognitive Sciences, MIT, Cambridge, MA; Laboratory of Neural Systems, The Rockefeller University, New York, NY; The Center for Brains, Minds, and Machines, MIT, Cambridge, MA

## Abstract

Vision must not only recognize and localize objects, but perform richer inferences about the underlying causes in the world that give rise to sensory data. How the brain performs these inferences remains unknown: Theoretical proposals based on inverting generative models (or “analysis-by-synthesis”) have a long history but their mechanistic implementations have typically been too slow to support online perception, and their mapping to neural circuits is unclear. Here we present a neurally plausible model for efficiently inverting generative models of images and test it as an account of one high-level visual capacity, the perception of faces. The model is based on a deep neural network that learns to invert a three-dimensional (3D) face graphics program in a single fast feedforward pass. It explains both human behavioral data and multiple levels of neural processing in non-human primates, as well as a classic illusion, the “hollow face” effect. The model fits qualitatively better than state-of-the-art computer vision models, and suggests an interpretable reverse-engineering account of how images are transformed into percepts in the ventral stream.

## Introduction

Perception confronts us with a basic puzzle: how can our experiences be so rich in content, so robust to environmental variation, and yet so fast to compute, all at the same time? Vision theorists have long argued that the brain must not only recognize and localize objects, but make inferences about the underlying causal structure of scenes^1–3^. When we see a chair or a tree, we perceive it not only as a member of one of those classes, but also as an individual instance with many fine-grained three-dimensional (3D) shape and surface details. These details can persist in long-term memory^4^ and are crucial for planning our actions – sitting in that chair or climbing that tree. Similarly, when seeing a face, we can not only identify a person if they are familiar, but also perceive so many details of shape, texture, and subtleties of expression even in people we have never met before.

To explain these inferences, early vision scientists proposed that scene analysis proceeds by inverting causal generative models, also known as “analysis-by-synthesis” or “inverse graphics”. Computational approaches to inverse graphics have been considered for decades in computational vision^3, 5–8^, and these models have some behavioral support^9, 10^. However, inference in these models has traditionally been based on top-down stochastic search algorithms, such as Markov Chain Monte Carlo (MCMC), which are highly iterative and implausibly slow. A single scene percept may take many iterations to compute via MCMC (which could be seconds or minutes on conventional hardware), in contrast to processing in the visual system which is nearly instantaneous. While top-down processing likely plays a role in some of the brain’s visual computations, such as surface segregation in complex scenes^11, 12^, both humans and non-human primates can extract rich high-level information about objects, faces and scene gists in a time window (150 milliseconds or less) that requires much (if not all) processing to be driven by a single feedforward pass^13–16^.

In part for these reasons, modern work in computational vision and neuroscience has focused on a different class of architectures, deep convolutional neural networks (DCNNs), which are more consistent with the fast, mostly feedforward dynamics of vision in the brain, and which benefit from simple, direct hypotheses about how their computations map onto neural circuits^13, 16^. DCNNs consist of many layers of features arranged in a feedforward hierarchy, typically trained discriminatively to optimize recognition of objects or object classes from labeled data. They have been instrumental both in leading engineering applications^17–19^ and in predicting neural responses in the primate visual system, both at the level of single units in macaque cortex as well as fMRI in humans^20–23^. Despite their impressive successes, however, conventional DCNNs do not attempt to address the question of how vision infers the causal structure underlying images. How we see so much so quickly, how our brains compute rich descriptions of scenes with detailed 3D shapes and surface appearances, in a few hundred milliseconds or less, remains a challenge.

Most recently, a new class of computational architectures has been developed that can potentially answer this challenge, by combining the best features of DCNNs and analysis-by-synthesis approaches. Several artificial intelligence (AI) research groups, including ours, have shown how neural network “inference models” can be built from a feedforward or recurrent network architecture trained to infer the underlying scene structure, rather than to recognize objects or classify object categories as in conventional DCNNs. In contrast to early analysis-by-synthesis algorithms, inference is fast, following a single bottom-up passes from the image or a small number of bottom-up-top-down cycles, without the need for extensive iterative processing^24–29^. These models have been developed in an engineering setting, and are just beginning to be tested in machine vision problems; their correspondence with human perception or neural mechanisms is unexplored. Here we introduce a specific model in this class, which we call the Efficient Inverse Graphics (EIG) network, and evaluate it as an account of face perception, arguably the best studied domain of high-level vision. The EIG model makes a number of fine-grained, quantitatively testable predictions, but also lets us evaluate the more general hypothesis that face perception in the brain is best understood in terms of an inference network that inverts a causal generative model (Fig. 1a), as opposed to the more conventional view in both AI and neuroscience that perception is best approached using neural networks optimized for classification, trained to recognize or distinguish object or face identities^16, 20, 30, 31^. We find that the EIG model is uniquely compatible with known data on human and non-human primate face processing, and provides the first quantitatively accurate and functionally explanatory account of both neural population responses in macaques and a range of challenging perceptual judgments in humans.

**Figure 1:**
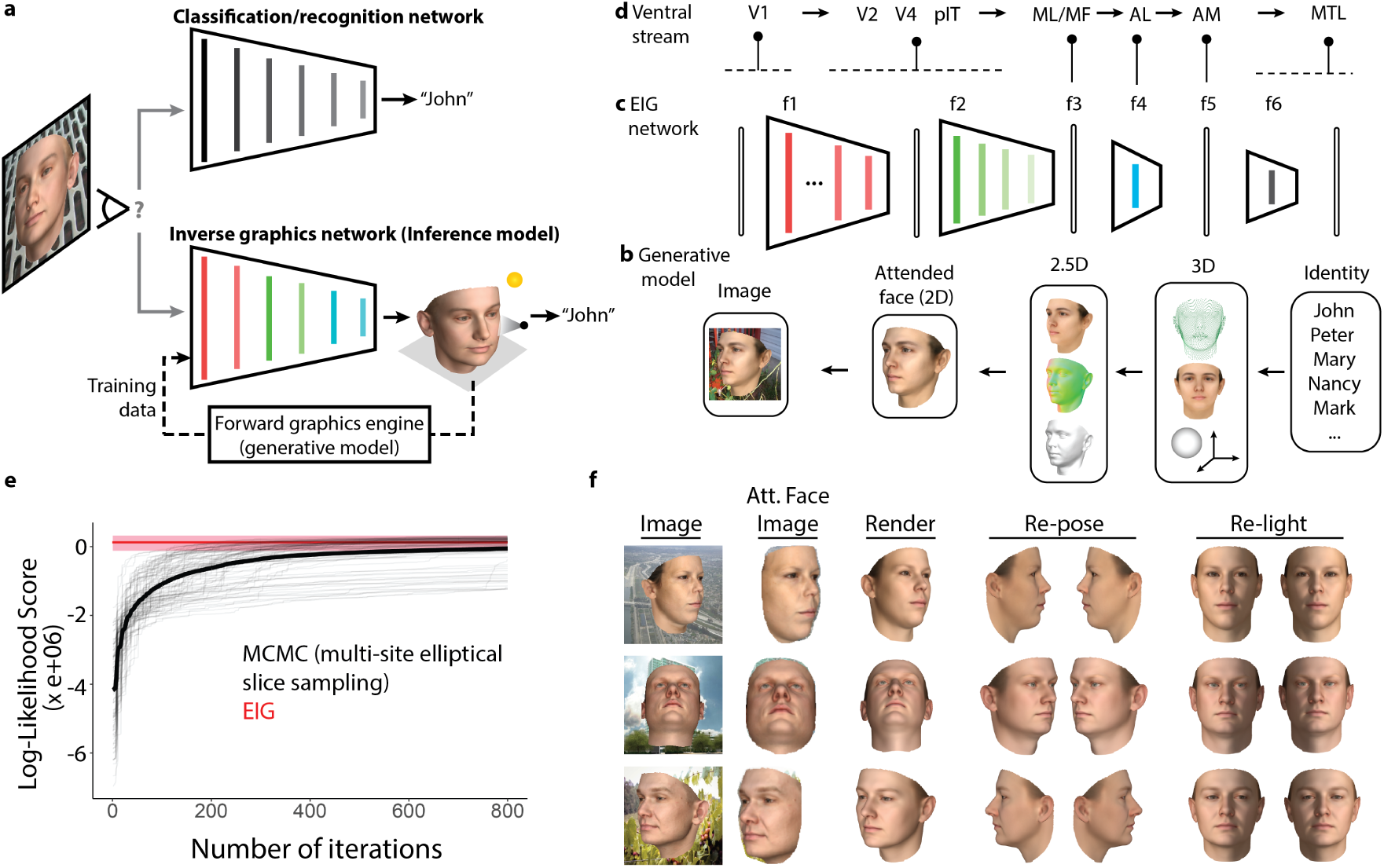
Overview of the modeling framework. **a**, Schematic illustration of two alternative hypotheses about the function of ventral stream processing: the recognition or classification hypothesis (top) and the inverse-graphics or inference network hypothesis (bottom). **b-d**, Schematic of the EIG model. Rounded rectangles indicate representations, arrows or trapezoids indicate causal transformations or inferential mappings between representations. **b**, The probabilistic generative model (right to left) draws an identity from a distribution over familiar and unfamiliar individuals, then through a series of graphics stages generates 3D shape, texture and viewing parameters, renders a 2D image via 2.5D image-based surface representations, and places the face image on an arbitrary background. **c**, The EIG inference network efficiently inverts this generative model using a cascade of DNNs with intermediate steps corresponding to intermediate stages in the graphics pipeline, including face segmentation and normalization (*f*_1_), inference of 3D scene properties via increasingly abstract image-based representations (convolution and pooling, *f*_2_ to *f*_3_) followed by two fully connected layers (*f*_4_ to *f*_5_), and finally a person identification network (*f*_6_). **d**, Schematic of ventral-stream face perception in the macaque brain, from V1 up through Inferotemporal cortex (IT), including three major IT face-selective sites (ML/MF, AL, AM), and on to downstream medial temporal lobe (MTL) areas where identity information is likely computed. Pins indicate empirically established or suggested functional explanations for different neural stages, based on the generative and inference models of EIG. Pins attached to horizontal dashed lines indicate untested but possible correspondences. **e**, Image-based log-likelihood scores for a random sample of observations using the EIG network’s inferred scene parameters (layer *f*_5_), compared to a conventional MCMC-based analysis-by-synthesis method. EIG estimates are computed with no iterations (red line; pink shows min-max interval), yet achieve a higher score and lower variance than MCMC, which requires hundreds of iterations to achieve a similar mean level of inference quality (thick line; thin lines show individual runs, see Methods). **f**, Example inference results from EIG, on held-out real face scans rendered against cluttered backgrounds. Inferred scene parameters are rendered, re-posed, and re-lit using the generative model.

The EIG model consists of two parts: a probabilistic generative model based on a multistage 3D graphics program for image synthesis (Fig. 1b), and an approximate inverse function of this generative model based on a DCNN that inverts (explicitly or implicitly) each successive stage of the graphics program (Fig. 1c), layer by layer. The inverse model, also known as an inference network or inference model, is the heart of EIG and the component we can most easily test with available neural and behavioral data. But the generative model is essential as well: it produces the training targets and training data for building the inference network, which is trained to infer the latent inputs or causes in the generative model conditioned on its outputs, rather than to predict class labels such as object categories or face identities as in conventional machine vision systems. The generative model, as we will see, also provides the basis for a functional interpretation of the representations the inference network learns. In this way, the EIG network embodies principles similar to the Helmholtz machine originally proposed by Hinton and colleagues in the 1990’s^32, 33^ and its modern cousins based on variational autoencoders (VAEs)^24, 25^. However, EIG differs from these approaches in that the generative model is based on an explicit graphics program (rather than a second deep neural network learned generically from data) and the EIG inference network is designed to parallel, in reverse, the graphics program’s structure. This allows EIG to more faithfully capture the causal processes of how real-world scenes give rise to images, and to exploit this structure for efficient learning and inference.

As a test case, we apply EIG in the domain of face perception where, in a rare co-occurrence, data from brain imaging, single-cell recordings, quantitative psychophysics and classic visual illusions all come together to strongly constrain possible models. EIG implements the hypothesis that the downstream targets of the ventral visual pathway, a series of interconnected cortical areas^34^ in inferotemporal (IT) cortex, are 3D scene properties analogous to the latent variables in a causal generative model of image formation (referred to as the “latent variables” or “inverse graphics” hypothesis; Fig. 1a); moreover, EIG specifies a precise circuit mechanism by which these properties are plausibly computed in the ventral stream (Fig. 1d). We compare EIG against a broad range of alternatives, including both lesions of EIG (leaving out components of the model) and multiple variants of state-of-the-art networks for face recognition in computer vision, implementing versions of the alternative hypothesis that the targets of ventral stream processing are points in an embedding space optimized for discriminating across facial identities (referred to as the “classification” or “recognition” hypothesis; Fig. 1a). We also consider alternative instantiations of the latent variables hypothesis, based on VAEs, which replace the structured generative graphics program in EIG with an unstructured generic deep neural network trained to reconstruct images. Only the EIG model, and therefore its more structured version of the latent variables hypothesis, accounts for the full set of neural and behavioral data, at the same time as it matches one of the most challenging perceptual functions of the ventral pathway: computing a rich, accurate percept of the intrinsic 3D shape and texture of a novel face from an observed image in a mostly feedforward pass.

### Efficient Inverse Graphics (EIG) Network

The core of EIG is the DCNN-based inference network, but we begin by describing the probabilistic generative model component, which determines the training objectives and produces the training data for the inference network. The generative model takes the form of a hierarchy of latent variables and causal relations between them representing multiple stages in a probabilistic graphics program for sampling face images (Fig. 1b). The top level random variable specifies an abstract person identity, *F*, drawn from a prior *Pr*(*F*) over a finite set of familiar individuals but allowing for the possibility of encountering a new, unfamiliar individual. The second level random variables specify scene properties: an intrinsic space of 3D face shape *S* and texture *T* descriptors drawn from the distribution Pr(*S, T |F*), as well as extrinsic scene attributes controlling the lighting direction, *L*, and viewing direction (or equivalently, the head pose), *P*, from the distribution Pr(*L, P*). We implement this stage using the Basel Face Model (a probabilistic 3D Morphable Model)^6, 35^, although other implementations are possible. These 3D scene parameters provide inputs to a z-buffer algorithm Ψ(·) that outputs the third level of random variables, corresponding to intermediate-stage graphics representations (or 2.5D components) for viewpoint-specific surface geometry (normal map, *N*) and color (albedo or reflectance map, *R*), {*N, R*} = Ψ(*S, T, P*). These view-based representations and the lighting direction then provide inputs to a renderer, Φ(·), that outputs an idealized face image, *I* = Φ(*N, R, L*). Finally, the idealized face image is subject to a set of image-level operations including translation, scaling, and background addition, Θ;(·), that outputs an observable raw image, *O* = Θ(*I*) (Fig. 1b; see Methods).

In principle, perception in this generative model can be formulated as MAP (Maximum A Posteriori) Bayesian inference as follows. We seek to infer the individual face *F*, as well as intrinsic and extrinsic scene properties *S, T, L, P* that maximize the posterior probability

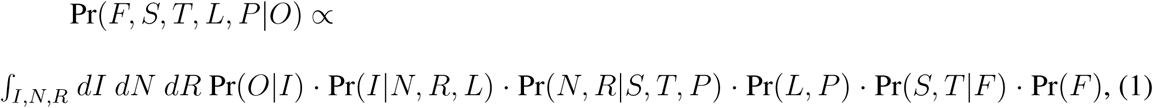

where Pr(*N, R|S, T, P*), Pr(*I|N, R, L*) and Pr(*O|I*) express likelihood terms induced by the mappings Ψ, Φ, and Θ respectively, and we have integrated out the intermediate representations of surface geometry and reflectance *N* and *R*, which perceivers do not normally have conscious access to, as well as the ideal face image *I*. Traditional analysis-by-synthesis methods seek to maximize Eq. 1 by stochastic local search, or to sample from the posterior by top-down MCMC inference methods; all of these computations can be very slow. Instead we consider a bottom-up feedforward inference model that is trained to directly estimate MAP values for the latent variables, *F *, S*, T *, L*, P \**.

This inference network (Fig. 1c) comprises a bottom-up hierarchy of functional mappings that parallels (in reverse) the top-down hierarchy of the generative model, and exploits the conditional independence structure inherent in the generative model for efficient modular inference. In general, if a random variable (or set of variables) *Z* renders two (sets of) variables *A* and *B* conditionally independent in the generative model, and if our goal is to infer *A* from observations of *B*, then an optimal (maximally accurate and efficient) feedforward inference network can be constructed in two stages that map *B* to *Z* and *Z* to *A* respectively^36, 37^. Here our inference model exploits two such crucial independence relations: (i) The observable raw image is conditionally independent of the 2.5D face components, given the ideal face image and (ii) The 2.5D components are conditionally independent of person identity, given the 3D scene parameters that describe the individual’s face. This conditional independence structure suggests an inference network with three main stages, which can be implemented in a sequence of deep neural networks where the output of each stage’s network is the input to the next stage’s network.

The first stage segments and normalizes the input image to compute the attended face image, the most probable value for the ideal image *I\** given the observed image *O*, by maximizing Pr(*I|O*) using a DCNN module trained for face volume segmentation^38^ and adapted to compute the face region given images of faces with background clutter (*f*_1_ in Fig. 1c).

The second stage is the core of our EIG model, and consists of a DCNN module trained to estimate intrinsic and extrinsic scene properties {*S∗, T ∗, L∗, P ∗*} maximizing Pr(*S, T, L, P |I∗*) from the attended face image. This network is adapted from the architecture of a standard “AlexNet” DCNN^17^ for object recognition, which consists of four convolutional layers (*f*_2_ in Fig. 1c) ending in a fifth, top convolutional feature space (TCL, *f*_3_ in Fig. 1c), followed by two fully connected layers (FCLs, *f*_4_ and *f*_5_ respectively). The training target for the second and final FCL *f*_5_ is the key difference from the conventional object recognition or face recognition pipeline: instead of being trained to predict class labels or identities, *f*_5_ is trained to predict scene properties, {*S, T, L, P* }. Training begins from a pretrained version of the basic architecture, fixing or finetuning weights up to layer *f*_4_, with only weights in the new scene property layer *f*_5_ being learned from random initial values. Training images for stage two are generated by forward-simulating images drawn from the generative model (in the spirit of the Helmholtz machine^32^), each with a different randomly drawn value for the scene parameters {*S, T, L, P* }, and using the generative model to produce the corresponding ideal face image *I* conditioned on those scene parameters.

Finally, a third inference stage estimates the most likely face identity label *F ∗* given the scene properties, maximizing Pr(*F |S∗, T ∗, L∗, P ∗*). Such identity labels are only introduced for familiar faces, with sufficient experience associating an individual’s identity to that face. This module comprises a single new FCL *f*_6_ for person identity classification, and is trained on labeled image-identity pairs. We generate these pairs from real-world experience if available, or by simulating real-world experience drawing faces randomly from the generative model and its prior over individuals *P* (*F*). In modeling particular experimental data, we tune training to the distribution of faces presented and introduce classification nodes for specific individuals if participants have sufficient opportunity to become familiar with them. See Methods section “EIG model” for further details of each of the three stages of the EIG network.

Together these three modules form a complete inference pipeline (approximately) inverting the generative model of face images, which satisfies the crucial characteristics of face perception and perceptual systems more generally: The inverse model (i) infers both rich 3D scene structure and the identities or class labels of individuals present in the scene, in a way that is robust to many dimensions of image variation and clutter, and (ii) computes these inferences in a fast, almost instantaneous manner given observed images.

We have tested the EIG inference network on both synthetic and held-out real face images, both isolated and superimposed on random backgrounds, and compared its performance with classic top-down analysis-by-synthesis algorithms based on MCMC^8^. EIG inferences are at least as accurate, assessed both quantitatively (Fig. 1e; see also Methods) and qualitatively (Fig. 1f), while being far faster (Fig. 1e). Thus EIG is at least a viable functional solution to the problem of face perception (but see Methods for a discussion of potential weaknesses as well). In the remainder of the paper, we ask how well the model captures the mechanisms of face perception in the mind and brain, by comparing its internal representations (especially *f*_3_, *f*_4_, *f*_5_) to neural representations of faces in the primate ventral stream, and its estimates of intrinsic and extrinsic face properties with the judgments of human observers in several hard perceptual tasks.

### Efficient inverse graphics stages explain macaque face-processing hierarchy

The best-understood neural architecture on which we can evaluate EIG as an account of perception in the brain is the macaque face-processing network^39^ [Fig. 2a; see Supplementary Information (SI) Section 4 for experimental procedure and neural recording details]. Freiwald and Tsao^39^ presented macaques with images of different individuals in different viewing poses (Fig. 2b), and found that this three-level hierarchy exhibits a systematic progression of tuning properties (Fig. 2c). Neurons in the bottom-level face patches ML/MF have responses driven largely by the pose of a face, independent of the face’s identity. Those in the mid-level patch AL also exhibit pose-specific tuning, but with a strong mirror-symmetry effect: faces in poses mirror-reflected about the frontal view axes exhibit similar responses. Neurons in the top-level patch AM exhibit view-robust identity coding. It has also been argued that these neural populations encode a multidimensional space for face, based on controlled sets of synthetically generated images.^40–42^. However, it remains unclear how the full range of three-dimensional shapes and appearances for natural faces viewed under widely varying natural viewing conditions might be encoded, and how high-level face space representations are computed from observed images through the multiple stages of the face-processing hierarchy.

**Figure 2:**
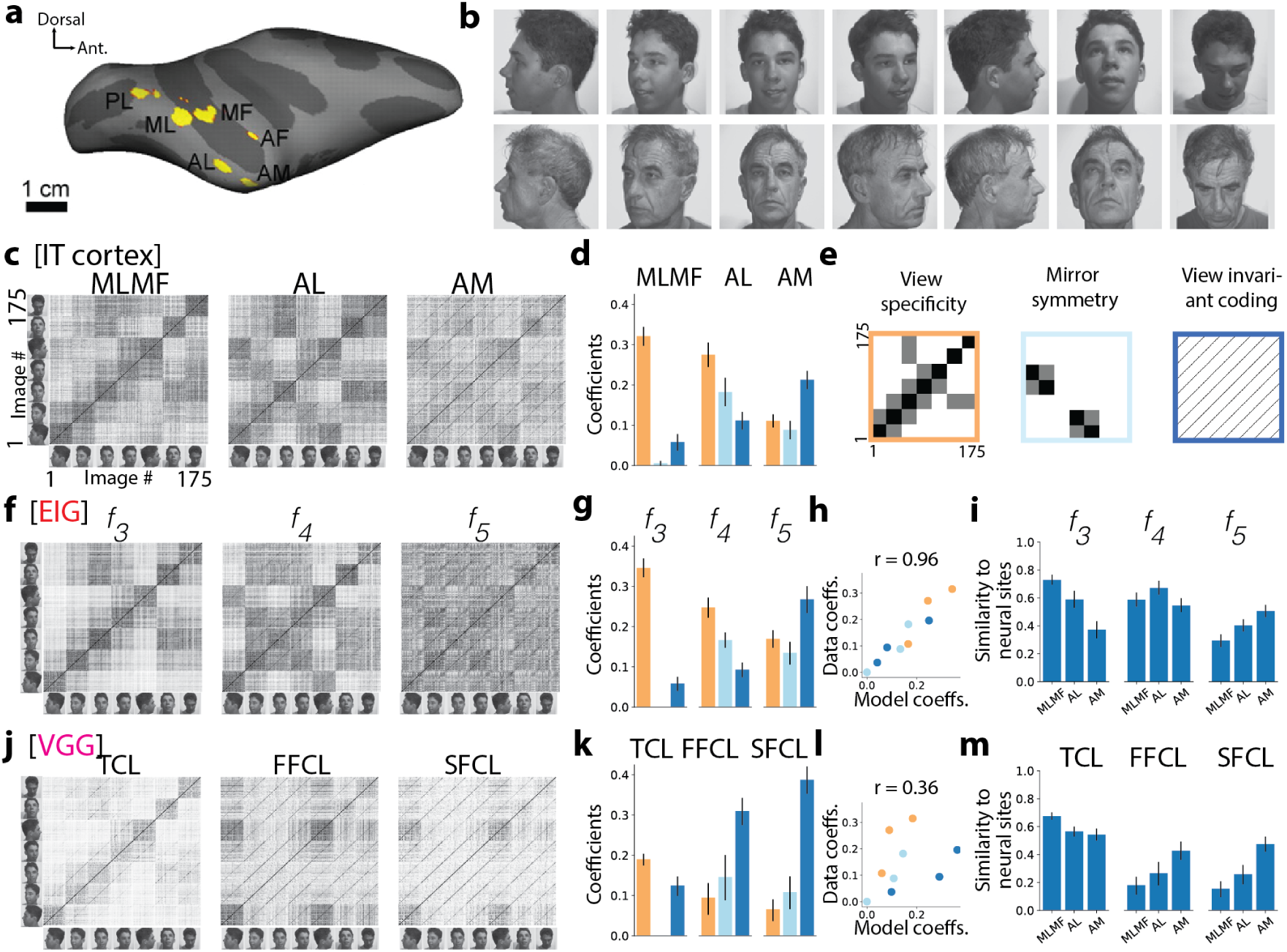
Inverse graphics in the brain. **a**, Inflated macaque right hemisphere showing six temporal pole face patches, including middle lateral and middle fundus areas ML/MF, anterior lateral area, AL, and anteriormedial area, AM. **b**, Sample FIV images. **c**, Population-level similarity matrices for each face patch. Each matrix shows correlation coefficients of population-level responses for each image pair from the FIV image set (25 individuals each shown in seven poses, total of 175 images)^39^. **d**, Coefficients resulting from a linear decomposition of the population similarity matrices in terms of idealized similarity matrices for view-specificity, mirror-symmetry, and view-invariance shown in (**e**), in addition to a constant background factor to account for overall mean similarity. **f**, Similarity matrices for each key layer of the EIG network, *f*_3_, *f*_4_, and *f*_5_, tested with FIV image set. Each image is represented as a vector of activations in the corresponding layer. **g**, Linear regression coefficients showing contribution of each idealized similarity matrix for each layer. **h**, Comparing full set of neural transformations to model transformations using these coefficients. **i**, Pearson’s r between similarity matrices of each neural population and model layer. **j-m**, VGG network tested using FIV image set. Panels follow the same convention as the EIG results. Error bars show 95% bootstrap confidence intervals (CIs; see Methods).

We address these questions by first quantifying the population-level tuning properties for the three successive levels of face patches, ML/MF, AL and AM, using linear combinations of three idealized similarity templates representing the abstract properties of view specificity, mirror symmetry, and view-invariant identity selectivity^43, 44^ (Fig. 2e) to fit the empirical similarity matrices for neural populations in each of these patches (see Methods). The coefficients of these different matrices (Fig. 2d) measure, in objective terms, how view-specificity decreases from ML/MF to AM (yellow bars), how mirror-symmetry peaks in AL (light blue bars), and how view-invariant identity coding increases from ML/MF to AL and further to AM (dark blue bars), complementing the qualitative features shown in the population-level similarity matrices (Fig. 2c).

We then evaluated the ability of the EIG network and other models to explain these qualitative and quantitative tuning properties of ML/MF, AL and AM. In particular we contrast EIG with several variants of the VGG network, based on a state-of-the-art DCNN for machine face recognition built via supervised training with millions of labeled face images from thousands of individual identities^31^ (see Methods). These comparisons allow us to tell apart the inverse graphics hypothesis and the classification hypothesis at the level of neural representation.

We first test models using the FIV set of natural face images, with 175 images of 25 individuals in 7 poses, shown to monkeys during neural recording of the face patches (Fig. 2b). The EIG network faithfully reproduces all patterns in the neural data, both qualitatively (Fig. 2f) and quantitatively in terms of the idealized similarity matrix analysis (Fig. 2g). The coefficients of all three idealized similarity templates (view specificity, mirror symmetry, view invariant coding) across all three levels of representation (*f*_3_/ML/MF, *f*_4_/AL, *f*_5_/AM) correlate almost perfectly between EIG and cortical face circuitry (*r* = 0.96, Fig. 2h). EIG also tracks the functional compartmentalization observed in the cortical hierarchy, as measured by raw correlations between similarities in corresponding layers: similarity in layer *f*_3_ best correlates with ML/MF, layer *f*_4_ best correlates with AL, and layer *f*_5_ best correlates with AM (*p <* 0.05, Fig. 2i). By all these measures, EIG appears to capture the full progression of three functionally distinct stages in face processing, from ML/MF through AL up to AM.

We evaluate VGG based on the three layers most analogous to EIG: the top convolutional layer (TCL), analogous to *f*_3_ of EIG, the first fully connected layer (FFCL) analogous to *f*_4_, and the second fully connected layer (SFCL) analogous to *f*_5_; these are also the layers of VGG most similar in response to the three levels of the face patch system. In contrast to EIG, VGG showed patterns of selectivity with some qualitative similarity to those of the neural data (Fig. 2j), but with pronounced qualitative and quantitative differences. Representations in VGG were substantially more view-invariant than either cortex or EIG across all three layers (*p <* 0.05, compare all yellow bars and dark blue bars in Fig. 2k versus 2d, g), with the biggest disparities occurring at the intermediate level (compare FFCL to *f*_4_ and AL in Fig. 2k, g, d). Indeed, there is no layer of VGG that shows the characteristic tuning of the intermediate patch AL, as *f*_4_ of EIG does, nor does any layer of VGG correlate maximally with AL relative to other neural sites; each layer is either a better fit to ML/MF or AM (Fig. 2m). Across all three levels in VGG, coefficients of the three idealized similarity matrices correlated much less strongly with analogous coefficients for neural data (*r* = 0.36, Fig. 2l), suggesting a failure to capture how face processing progresses through the cortical hierarchy. Most dramatically, the two highest layers of VGG (FFCL and SFCL) were almost indistinguishable from each other (Fig. 2j), which fails to reflect the clear progression in function from mirror-symmetric tuning to view invariant coding that is seen in both the corresponding layers of EIG (*f*_3_ and *f*_4_) and the corresponding neural sites (AL and AM).

Other analyses show that VGG performance does not depend on whether it is fine-tuned to these specific face identities (as in Fig. 2b; see Extended Data Fig. 2 for VGG in its raw pretrained state), and that the initial face segmentation and normalization stage of EIG, which has not been a component of previous ventral stream models^13, 16, 20^, is necessary for its strong performance (but has little effect on VGG; see Methods and Extended Data Fig. 2). Taken together, these results strongly support the hypothesis that ventral stream face processing begins with an initial segmenting operation and culminates in targets that encode the latent variables of a face generative model, rather than mapping raw images to features optimized for face identity recognition or discrimination, as in conventional machine vision approaches.

To better understand the reasons why a fully brain-like pattern of responses arises in EIG, and the conditions under which it might arise in other neural network models, we studied a large number of model alternatives, varying in network architecture, training set and objective, and standard aspects of training procedure (see Methods and SI Sections 1,2; Extended Data Figs. 4-7). We used a controlled synthetic analog of the FIV image set, in which only faces were rendered (without clothing or backgrounds; FIV-S; see Methods). In particular, we tested several VAE variants that shared EIG’s feedforward inference-network architecture but used a different training objective (image reconstruction loss; Extended Data Fig. 5) and used deep neural networks to parametrize a learned generative model (as opposed to EIG’s structured graphics engine). We also tested several variants of the VGG architecture (Extended Data Fig. 4) to unconfound effects of the VGG architecture, training set and training objective. We say that a model produces a “fully brain-like pattern of responses” to the extent that it has three progressive layers with idealized similarity coefficients matching those in ML/MF, AL, AM (i.e., the bar plots shown in Fig. 2d), correlating highly across layers (as in Fig. 2h), and with raw similarities in each of these model layers correlating maximally and distinctively with raw similarities in the corresponding neural sites (as in Fig. 2i). Two aspects of the EIG network, its training targets and architecture, proved necessary to obtain fully-brain like representations: (1) The targets of inference should be the latent variables of the causal generative model (3D face shape and face texture descriptors), and (2) There should be a stack of convolutional layers processing the attended face image followed by at least one fully connected hidden layer between the top convolutional layer and the final layer trained to estimate the latent variables. Other aspects of the EIG training procedure, such as the magnitude of dropout and initialization with pretrained network weights, were not essential for producing fully brain-like responses but do make training much more efficient (Extended Data Figs. 6,7).

Finally we ask whether intermediate stages of the face-processing hierarchy, ML/MF and AL in the primate brain or *f*_3_ and *f*_4_ in the EIG network, can be given an interpretable functional account as we did for AM and *f*_5_, or whether instead these patches are best understood simply as a hierarchy of “black box” function approximators. Fig. 1b and c suggest one possible functional interpretation based on correspondences between the graphics and inverse-graphics pathways: ML/MF could be understood as computing a reconstruction of an intermediate stage of the generative model, the 2.5D components of a face (e.g., albedos and surface normals, or surface depths) analogous to the “intrinsic images” or “2.5D sketch” of classic computer vision systems^3, 45^. It is also possible that these patches compute a reconstruction of an earlier stage in the generative model such as the attended face image (corresponding to the output of *f*_1_), or that they are just stepping stones to higher-level representations without distinct functional interpretations in terms of the generative graphics model. We computed similarity matrices for each of these candidate interpretations (each generative model stage), as well as for the raw pixel images as a control (Fig. 3a; see SI Section 3 for how 2.5D components of the FIV images are approximated). We then correlated these similarity matrices with those for ML/MF and AL. We find that the 2.5D components best explain ML/MF (*p <* 0.001), and closely resemble their overall similarity structure (Fig. 3b). Attended images also provide a better account of ML/MF than the raw pixel images (*p <* 0.001) but significantly worse than the 2.5D components (*p <* 0.001 for each component; Fig. 3b). We also find that the 2.5D components explain *f*_3_ layer responses in the EIG model better than the raw pixel images, and better than the attended face image when these can be discriminated (see SI section 3; Extended Data Fig. 8).

**Figure 3:**
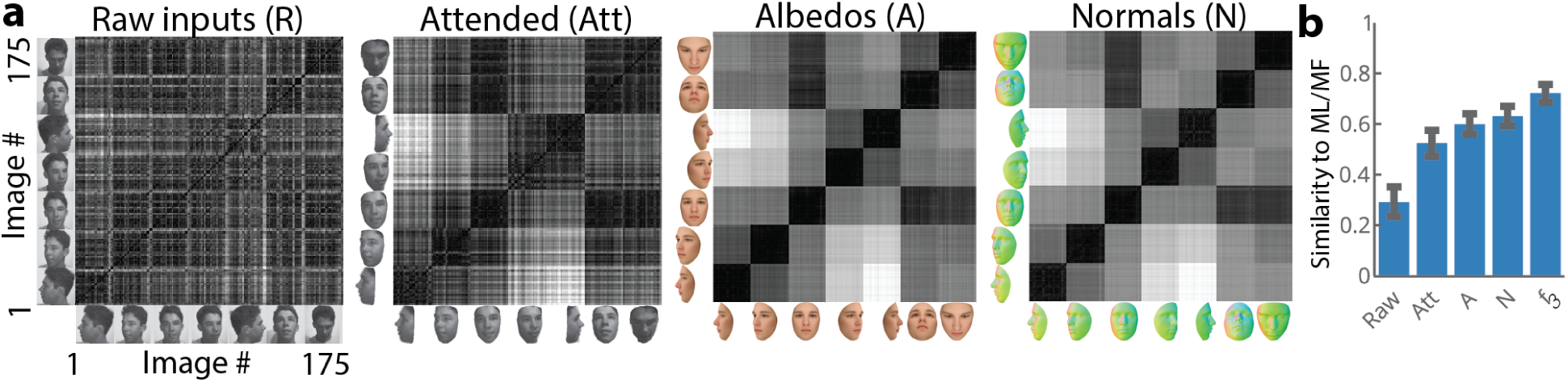
Understanding ML/MF computations using the generative model and the 2.5D (or intrinsic image) components. **a**, Similarity matrices based on raw input images, attended images, albedos, and normals. Colors indicate the direction of the normal of the underlying 3D surface at each pixel location. **b**, Correlation coefficients between ML/MF and the similarity matrices of each image representation in (**a**) and *f*_3_. Error bars indicate 95% bootstrap CIs.

AL has no such straightforward representational account, but it may be understood as implementing a densely connected hidden-layer mapping the estimated 2.5D face components (in ML/MF and *f*_3_) to estimated 3D face properties (in AM and *f*_5_). This highly nonlinear transformation can be facilitated using some kind of hidden layer, and this could be the role of AL in the primate brain and the corresponding layer *f*_4_ in EIG. Note that such an intermediate layer appears to be functionally missing from VGG and its variants trained to discriminatively predict identity rather than 3D object properties. These models always show very similar responses in all their fully connected layers (compare Fig. 2c and 2j and also see Extended Data Fig. 4). We conjecture that this AL-like intermediate stage nonlinearity is not necessary because the fully connected layers of VGG are solving a different task than EIG or the brain: VGG appears to be mapping high-level image features (computed at the top of the convolutional layers) to person identities which are almost linearly decodable from these features, without ever having to explicitly represent the 3D properties of a face (see SI Section 3; Extended Data Fig. 9). The VGG network design may be a reasonable, perhaps even superior, way to build a system for face perception if the goal is merely to classify or recognize individuals through their facial appearance, as in most of today’s computer vision system. But the brain needs to compute much richer information about the 3D shape and texture of faces, in order to analyze expressions, emotions, mood and health, or to use face perception as a cue in spoken language understanding. The inverse graphics design of the EIG network offers a possible route to those richer percepts, and our analyses suggest that the ML/MF-AL-AM circuit may be the locus of these computations in the brain.

### Efficient inverse graphics scene parameters predict human behavior

We also tested EIG and alternative models’ ability to explain the behavioral aspects of face perception, by comparing their responses to people’s judgments in a suite of challenging unfamiliar face recognition tasks^46^. In three experiments (inspired by the passport photo verification task), subjects were asked to judge whether two sequentially presented face images showed the same or different identity (Fig. 4a). In Experiment 1 (“Regular”), both study and test images were presented with pose and lighting directions chosen randomly over the full range covered by the generative model. Experiments 2 and 3 probed generalization abilities, using the same study items from Experiment 1 but test items that extended qualitatively the range of training stimuli. In Experiment 2 (“Sculpture”), the test items were images of face sculptures (i.e., texture-less face shapes rendered with a stone-like uniform grey albedo in frontal pose) eliminating all cues from skin coloration or texture normally present in face inputs. In Experiment 3, the test items were flat frontal facial textures, produced by distorting normal images using a fish-eye lens effect to reduce shape information in the input (see Methods).

**Figure 4:**
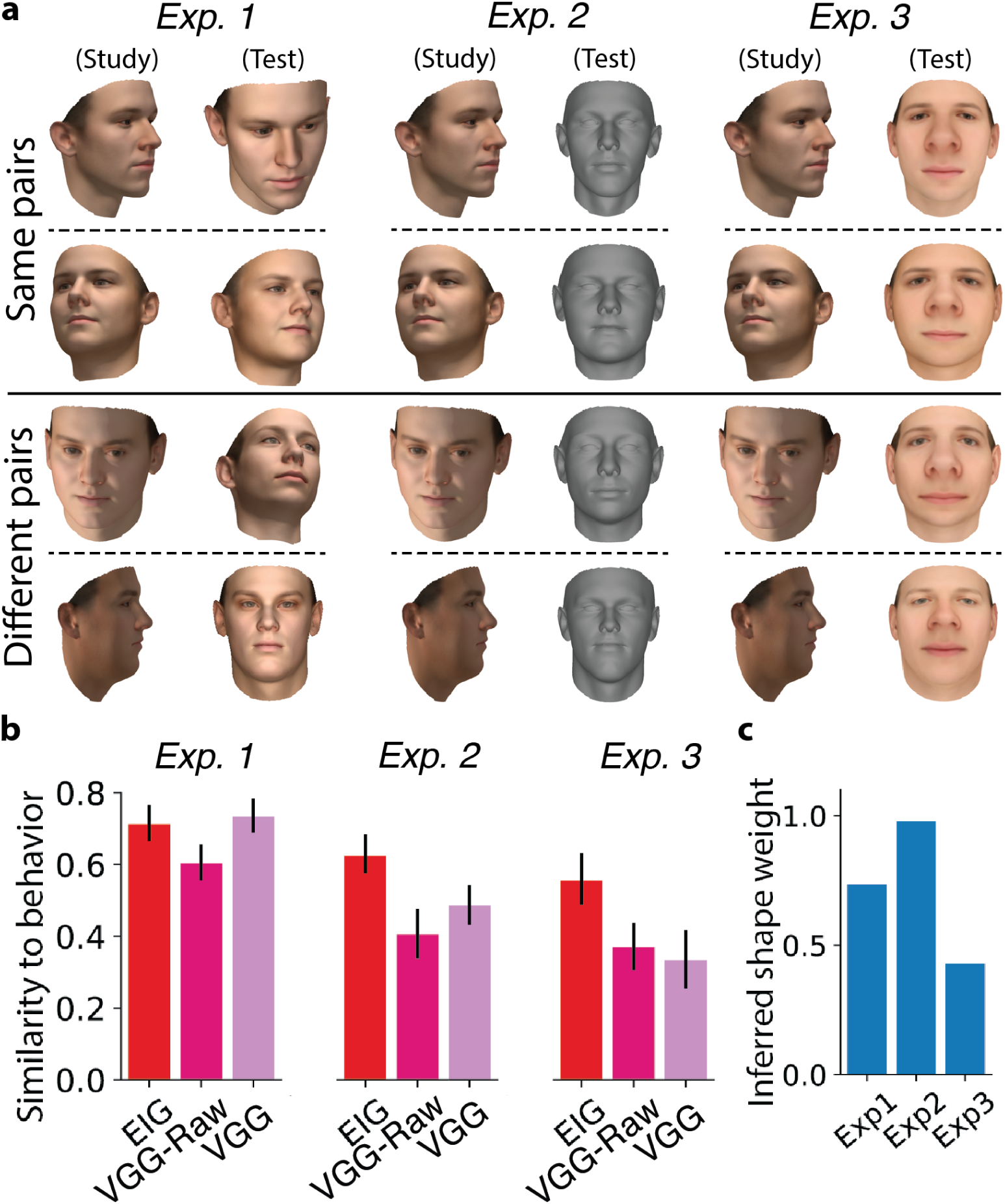
Across three behavioral experiments, EIG consistently predicts human face recognition performance. **a**, Example stimuli testing same-different judgments (“same” trials rows 1-2, “different” trials rows 3-4) with normal test faces (Exp. 1), “sculpture” (texture-less) test faces (Exp. 2), and fish-eye lens distorted shade-less facial textures as test faces (Exp. 3). **b**, Correlations between model similarity judgments and human judges’ probability of responding “same”. **c**, Inferred weights (a value between 0 and 1 that maximized model’s recognition accuracy) of the shape properties (relative to texture properties) in the EIG model predictions for Exps. 1-3. Error bars indicate 95% bootstrap CIs (see SI Section 5).

We hypothesized that if face perception is based on inverting a generative model with independent 3D shape and texture latents, as in EIG but not VGG, VGG-Raw or other classification/recognition alternatives, then participants might be able to selectively attend to shape or texture estimates in their internal representations, in order to optimize performance on these different challenge tasks. Crucially, EIG and VGG models are both trained using an equal number of images synthesized from the same graphics program used to generate the stimuli (although VGG is finetuned on top of the VGG network which itself is trained with millions of other face images); only their training targets are different: latent variables of the generative face model for EIG, versus an embedding space for discriminating person identities for VGG. This allows our behavioral analyses, like our neural analyses, to test between the two different hypotheses about the functional goal of face perception, inference in a generative model versus classification or recognition of individuals’ identities.

For each experiment, we compared average human responses – i.e., Pr(“Same”), frequency of the “same” response – to the models’ predicted similarity across trials. A model’s predicted similarity for a given trial was computed as the similarity between the model’s outputs (i.e., its top layer) for the study and test items (see Methods). The VGG and VGG-Raw networks’ outputs for an image is their identity-embedding spaces, the layer SFCL. (No other layer in the VGG network provided a better account of the human behavior than its SSFL layer.) EIG’s output is its shape and texture parameters, which unlike other models supports selective attention to these different aspects of a face. For each experiment we fit a single weight for the shape parameters in EIG’s computation of face similarity (constant across all trials and participants); the weight of the texture component is 1 minus that value.

Overall, participants performed significantly better than chance (50% correct): Average performance was 66% correct in Experiment 1, 64% in Experiment 2, and 61% in Experiment 3. (See SI Section 5 for model-free behavioral analysis.) In trial-by-trial comparisons to behavior, EIG consistently predicted human error patterns across all three experiments, with *r* values 0.70[0.65, 0.76], 0.64[0.58, 0.69], and 0.54[0.47, 0.61] (where [*l, u*] indicates lower/upper 95% confidence intervals; Fig. 4b). VGG (though not VGG-Raw) performed comparably on Experiment 1, but EIG fit human judgments significantly better than both alternative models in Exps. 2 and 3 (*p <* 0.001 for all comparisons based on direct bootstrap hypothesis tests; see SI Section 5). EIG’s inferred attention weight showed a bias towards shape properties in Experiment 1 and a bias towards texture properties in Experiment 3, but it attended to shape parameters almost exclusively for Experiment 2 (inferred weight 0.99; Fig. 4c). These results suggest that EIG captures human face perception abilities more accurately than other models, especially under less familiar stimulus conditions and tasks requiring extreme generalization between study and test faces. They also lend further support to the latent variables hypothesis over the classification hypothesis for ventral stream face processing.

Human face perception is susceptible to illusions, and our model naturally captures one of the most famous. In the hollow face illusion, a face mask reversed in depth (so the nose points away from the viewer) appears to be a normally shaped face with two distinctions: (i) hollow faces lit from the top or side appear to be lit from the bottom or alternate side^47^, and (ii) hollow faces appear flatter than normal faces^48^. It has been suggested that this illusion could be a result of Bayesian inference, arising from the integration of top-down priors for natural face geometry, appearance and lighting with ambiguous bottom-up cues to depth such as shading patterns^48, 49^. To our knowledge, this proposal has not previously been tested quantitatively or implemented in a working computational model. Here, we psychophysically study the hollow-face effect in greater detail using graded levels of depth reversals, and test EIG quantitatively as a computational account of human illusory percepts at a trial-level granularity.

We compared our model’s inferences about lighting direction and face depth with people’s judgments, in both graded versions of the hollow face illusion and normal lighting direction variation, as a control (Fig. 5a, b). We found that the EIG network, like humans, perceived the light source direction to covary illusorily with graded reversal of the face depth map, in a highly nonlinear pattern inflecting just when depth values turned negative; in contrast, varying lighting direction in a normal way while keeping face shape constant (the control condition) was perceived linearly and largely veridically by both people and the model (Fig. 5c,d). We also found that the EIG network, like humans, perceived depth-inverted faces as more flat when compared to their control counterparts with the lighting source elevation matched to its illusorily perceived location in the depth-inverted condition; indeed, the EIG network closely matched the magnitude of flattening in depth judgments as a function of the level of depth reversal for hollow faces, as well as a subtle effect of lighting elevation on judged depth in the control condition (Fig. 5e,f). We also attempted to decode these same lighting and profile depth parameters from the VGG network, and found significantly worse fits to human judgments in all cases, but especially in depth judgments where VGG fits were barely better than chance (see SI Section 5; Extended Data Fig. 13). The fact that the EIG network captures the nonlinear interaction of depth and lighting percepts in the hollow face illusion does not uniquely support EIG as an account of the ventral face pathway; a “vanilla” network could be trained to estimate either lighting or profile depth from face images and might predict the same judgments. Rather, EIG’s success here relative to VGG, without EIG having to be trained specially on these atypical images or ever being trained explicitly to estimate profile depth, provides further evidence that ventral stream face perception as modeled by EIG is implementing some form of fast approximate analysis-by-synthesis or inverse graphics computation, as opposed to being optimized for recognition of face identity.

**Figure 5:**
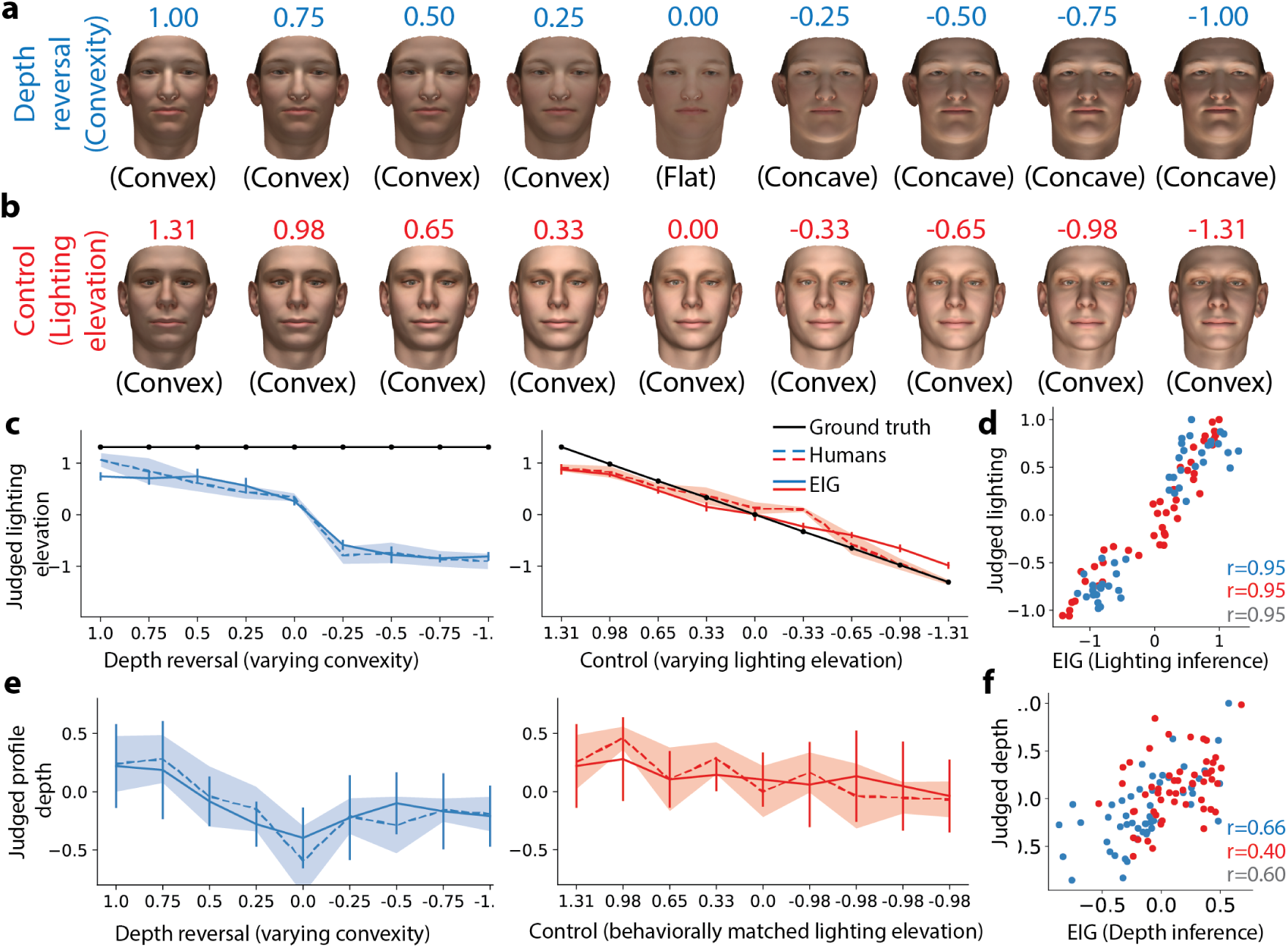
Psychophysics of the “hollow face” effect. On a given trial, participants saw an image of a face lit by a single light source and judged either the elevation of the light source (**c**, **d**) or the profile depth of the presented face (**e**, **f**) using a scale between 1 and 7 (also see Methods). **a**, One group of participants (depth suppression group) were presented with images of faces that were always lit from the top, but where the shape of the face was gradually reversed from a normally shaped face (convexity=1) to a flat surface (convexity=0) to an inverted hollow face (convexity=-1). **b**, Another group of participants (control group) were presented with images of normally shaped faces (convexity=1) lit from one of the 9 possible elevations ranging from the top of the face to the bottom. **c**, Normalized average light source elevation judgments of the depth-suppression group (left), the control group (right), EIG’s lighting elevation inferences, and the ground truth light source location. **d**, Average human judgments vs. EIG’s lighting source elevation inferences across all 90 trials without pooling to 9 bins. Pearson’s r values are shown for all trials (grey), control trials (red), and depth suppression trials (blue). **e**, Normalized average profile depth judgments of the depth-suppression group (left), control group (right) and EIG’s inferred profile depth. **f**, Average human judgments vs. EIG’s inferred profile depths across all 108 trials without pooling to 9 bins. Pearson’s r values are shown as in (**d**).

## Discussion

Our results suggest that the primate ventral stream approaches at least the first feedforward pass in face perception – and perhaps object perception more generally – with an “inverse graphics” strategy implemented via an efficient hierarchical inference network: Observed images are mapped via a segmentation and normalization mechanism to a 2.5D-like map of intrinsic surface properties (view-centered geometry and albedo) represented in ML/MF, which is then mapped via a nonlinear transform through AL to a largely viewpoint-independent representation of 3D object properties (shape and texture) in AM. The EIG network simulates this process and captures the key qualitative and quantitative features of neural responses across the face-patch system, as well as human perception for both typical and atypical face stimuli. The EIG model thus suggests how the structure of the visual system might be optimized for its function: computing a rich representation of behaviorally relevant causal properties underlying the appearance of a novel object or scene, as quickly and as accurately as possible.

Our results are consistent with strong evidence that neurons in areas ML/MF and AM code faces in terms of a continuous “shape-appearance” space^42^, not simply discrete identities. However, the EIG model goes beyond this finding to address core, long-standing questions of neural computation: How is the ultimate percept of an object (or face) derived from an image via a hierarchy of intermediate processing stages, and why does this hierarchy have the structure it does? EIG is an image-computable model that faithfully reproduces representations in all three face patches of ML/MF, AL and AM, and explains mechanistically how each stage is computed. It also suggests why these representations would be computed in the sequence observed, in terms of a network for moving from 2D images to 2.5D surface components to 3D object properties, which exploits the conditional independence properties of a generative model for how face scenes produce images to efficiently invert that process. The model thus gives a systems-level functional understanding of perhaps the best characterized circuitry in the higher ventral stream.

Anatomical connectivity and temporal dynamics of responses in the face patches suggest the existence of feedback and other non-hierarchical connectivity that our current model does not capture^50^. Following earlier models of primate face and object processing^16, 20, 51^, we see a feed-forward hierarchical network such as EIG as only a first approximation of the system’s functional architecture – a natural starting point, as so much rich information about faces (and objects and scenes) is already computed in the first 150 msecs of feedforward inference, but clearly just a first step that future work should go beyond. More generally, there are important functions of vision that can be understood in terms of inverting generative models, such as segregating multiple objects or surfaces in complex or cluttered scenes, which appear to depend on feedback or recurrent connections, especially to early visual areas (V1/V2)^11, 12^. Explaining these neural computations could benefit greatly from the study of efficient inverse graphics architectures that integrate bottom-up and top-down processing^28, 52^. It is also possible that such feedback architectures could provide a fuller account of the mechanisms by which the computations in our EIG network are implemented in the brain.

That our model simultaneously explains the full macaque face patch system and the outputs of human psychophysical judgment provides further support that human and non-human primate face systems share similar organization^53, 54^. Recent work comparing VGG face network representations with neural representations in humans using intracranial EEG (iEEG) data^30^, taken together with our results here, also suggests a consistent picture. Grossman et al. find evidence that VGG only matches human face representations up to the model network’s TCL, in areas of human IT thought to best correspond to the middle face patches we study here (ML/MF). They take this as evidence that human face circuitry is performing a more “pictorial” form of processing than VGG’s recognition computations, but they do not specify an alternative network architecture or concrete computational hypothesis for what that pictorial processing might be. Our model suggests one such hypothesis, in the form of 2.5D or intrinsic image components, which capture facial appearance and shape in a view-based, image-centric frame, and correspond well to middle face patch representations in macaques. Our model also suggests how those pictorial 2.5D representations can lead downstream to a full 3D description of face shape and appearance, which would correspond to more anterior face regions that (as Grossman et al. note) have yet to be studied intracranially in humans, and then further downstream to representations of familiar individuals’ identies (e.g, MTL, perirhinal cortex), which have been well-characterized in both humans^55^ and macaques^56^.

Our approach also has broader implications for neuroscience, perception, and cognition. The finding that IT supports decoding of category-orthogonal shape information for a wide range of objects, in addition to object category identity^57^, suggests that an extension of EIG could account for how the brain perceives the three-dimensional structure of objects beyond the domain of faces. With other collaborators, we have recently shown in an AI context that EIG-like networks for efficient inference of 3D shapes from 2D images via 2.5D sketches can work for arbitrary object classes (e.g., chairs, cars)^29, 58^, and can even generalize to a range of novel, unseen classes^59^. In future work, we hope to explore these models of how the ventral visual pathway processes other object classes with functionally specific, localized representations (bodies, hands, word forms), as well as objects more generally.

If this larger program is successful, it may offer a resolution to the problem of interpretability in visual neuroscience^60^: Today’s best performing models are remarkable for their ability to fit stimulus-dependent variance in neural firing rates, but often without an interpretable explanation of what those neurons are computing. Our work suggests that in addition to maximizing variance explained, computational neuroscientists could aim for “semi-interpretable” models of perception, in which some neural populations (such as ML/MF and AM) can be understood as representing stages in the inverse of a generative model (such as 2.5D components and 3D shape and texture properties), while other populations (such as AL) might be better explained as implementing necessary hidden-layer (nonlinear) transforms between interpretable stages.

The efficient inverse graphics approach can also be extended to richer perceptual inferences where there is currently no consensus on how these computations are implemented in the brain. EIG networks can be augmented with multiple scene layers in order to parse faces or other objects under occlusion^61, 62^. They be deployed in parallel or in series (using attention) to parse out multiple objects in a scene^24, 63–66^. They can even be extended to other modalities through which we perceive physical objects, such as touch, and can support flexible crossmodal transfer, allowing objects that have only been experienced in one modality (e.g., by sight) to be recognized in another (touch)^61^. All of these extensions suggest testable hypothesis for neural computations and representations, in ways that could also point to crucial functional roles for feedback or recurrent processing, which our work here does not address.

Finally, while our work suggests a functional role for causal generative models in the visual system, it leaves open many questions about their nature, use, and origins. Interpreted most literally, EIG implies that the brain uses feedforward inference networks as the workhorse of object perception, but uses generative models to provide the targets for training those networks, and as a source of internally generated training data (possibly at multiple stages, in a recognition pipeline that inverts a multi-stage generative process). Generative models in the brain could also support other functional roles, however: They could be used during online perception to refine a percept – particularly in hard cases such as under dim light or under heavy occlusion– by enforcing reprojection consistency with intrinsic image based surface representations^8, 26, 29, 63^. They could also support higher functions in cognition such as mental imagery, planning, and problem solving^67, 68^. It remains to be determined which of these functions are actually operative in the brain, as well as where and how generative models might be implemented in neural circuits, and how they might be built over development, from some combination of genetically programmed mechanisms and early perceptual experience. VAEs, and their close cousins GANs^69^, capsules^52^, and GQNs^24^, as well as RCNs^28^, are recent developments in artificial network architectures that suggest at least partial hypotheses for how graphics models might be implemented neurally, or constructed through learning, but none of these suggestions are yet well grounded in experimental work. We hope that the success of the EIG approach here will inspire future work to explore potential neural correlates of these architectures, as well as the other roles that generative models could play in perception, cognition, and learning.

## Methods

### Generative model

Our generative model builds on and extends the Basel Face Model (BFM)^1^, a statistical shape and texture model obtained by applying probabilistic principal component analysis^2^ on a dataset of 200 laser-scanned human heads (100 female, 100 male). BFM is publicly available and consists of a mean (or norm) face shape, a mean texture, two sets of principal components of variance, one for shape and the other for texture, and their corresponding eigenvectors that projects these principal components to 3D meshes.

The principal components of shape *S* and texture *T* accept a standard normal distribution such that Pr(*S*) and Pr(*T*) are each multivariate standard normal distributions with 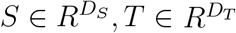. Each sample from Pr(*S*) (or Pr(*T*)) is a vector in a *D* = *D*_*S*_ (or *D* = *D*_*T*_) dimensional space specifying a direction and a magnitude to perturb the mean face shape (or the mean texture) to obtain a new unique shape (or texture). Mean shape and texture correspond to *s* = {0, 0,*…*, 0} and *t* = {0, 0,*…*, 0}. (Uppercase letters are used for random variables and lowercase letters are used for assignments of these random variables to a sample from their respective distributions. Nonrandom model parameters, such as *D* are also uppercase.) We set *D*_*S*_, *D*_*T*_ = 200 in our analysis. We found that the exact values of *D*_*S*_ and *D*_*T*_ did not matter as long as they were not too small, which leads to very little variation across the samples.

We used the part-based version of BFM where the principal components of shape and texture are partitioned across four canonical face parts: (i) outline of the face, (ii) eyes area, (iii) nose area, and (iv) mouth area. Each face-part (e.g., shape of the nose area or texture of the eyes area, etc.) was represented using 200*/*4 = 50 principal components. There are four advantages of using BFM: it (i) allows a separable representation of shape and texture, (ii) provides a probability distribution over both of these properties, (iii) allows us to work with lower dimensional continuous vectors (400 dimensions in this case) as opposed to very high dimensional meshes (e.g., meshes consisting of about 1 million vertices), and (iv) consists of dimensions that are often (but not always) perceptually interpretable (e.g., a dimension controlling the inter-eye distance).

The full scene description in the model also requires choosing extrinsic scene parameters including the lighting direction and viewing direction (or equivalently, head pose). In our simulations, we used Lambertian lighting where the lighting direction *L* can vary along azimuth *L*_*a*_ and elevation *L*_*e*_. Pr(*L*_*a*_) and Pr(*L*_*e*_) are uniform distributions in the range {-1.4^*rad*^} to {1.4^*rad*^}. The head pose *P* can vary along the z-axis *P*_*z*_ with Pr(*P*_*z*_) a uniform distribution in the range −1.5^*rad*^ to 1.5^*rad*^, and the x-axis *P*_*x*_ with Pr(*P*_*x*_) a uniform distribution in the range −0.75^*rad*^ to 0.75^*rad*^. Finally, we rendered each scene to a 227 × 227 pixels color image, unless otherwise mentioned, with back-face culling.

#### Synthetic FIV image sets

The FIV-S stimuli underlying Extended Data Fig. 4-8 used the pose distributions in Extended Data Table 1. Each of the 25 identities (i.e., unique pairs of shape and texture properties) were rendered at 7 different poses and with frontal lighting.

The image set underlying Extended Data Fig. 8B (referred to as FIV-S-2) used the same prior over lighting and pose as the generative model, Pr(*L*) and Pr(*P*), described above. It used the same 25 identities as FIV-S image set each rendered 7 times (each with its own randomly drawn pose and lighting parameters) making 175 images in total. Additionally, to increase the variability at the level of raw and attended images, we converted half of these images to gray-scale.

#### Conventional top-down inference with MCMC

Given a single image of a face as observation, *I*, and an approximate rendering engine, *G*(·) – a combination of the z-buffer Ψ(·) and image rendering Φ(·) stages introduced in the main text – face processing in this probabilistic graphics program can be defined as inverting the graphics pipeline using Bayes’s rule:

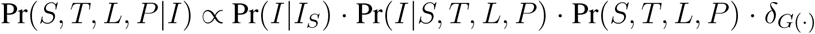

where I_*S*_ is a top-down sample generated using the probabilistic graphics program, and *δ*(·) is a Dirac delta function. (We dropped the corresponding Dirac delta functions in Equation 1 in the main text in order to avoid cluttered notation.) We assume that the image likelihood is an isotropic standard Gaussian distribution, *P* (*I|I*_*S*_) = *N* (*I*; *I*_*S*_, Σ). Note that the posterior space is of high-dimensionality consisting of more than 400 (404, to be exact) highly coupled shape, texture, lighting direction, and head pose variables, making inference a significant challenge.

Markov chain Monte Carlo (MCMC) methods provide a general framework for inference in generative models and have a long history of application to inverse graphics problems^3^. For this specific face model, we explored both traditional single-site MCMC and a more advanced and efficient multi-site elliptical slice sampler^4^ to infer the shape and texture properties given an image, *I*_*D*_^5^. Proposals in elliptical slice sampling are based on defining an ellipse using an auxiliary random variable *X ∼ N* (0, Σ) around the current state of the latent variables (shape and texture properties), and sampling from an adaptive bracket on this ellipse based on the log-likelihood function. For the lighting direction and pose parameters, single-site Metropolis-Hastings steps are used. At each MCMC sweep, the algorithm iterates a proposal-and-acceptance loop over twelve groups of random variables: four shape vectors (each of length 50), four texture vectors (each of length 50), and four scalars for lighting direction and pose parameters. The detailed form of the proposal and acceptance functions can be found in^4^. This method often converges to reasonable inferences within a few hundred iterations, although with substantial variance across multiple runs of the algorithm as shown in Fig. 1c in the main text. The y-axis values in that figure are the log-likelihood scores *P* (*I|S, T, L, P*) of 100 individual chains each given as input a different face image (with clean background). The log-likelihood score for each iteration of each chain is calculated by rendering and comparing the current MCMC estimate with the input image. The log-likelihood scores for the EIG network on Fig. 1c are computed in the same way except its estimates are outputs at its layer *f*_5_.

The EIG estimates are computed almost instantaneously, with no iterations, yet achieve a higher score and lower variance (mean score, red line, *∼* 2.5 *×* 10^5^; standard deviation *∼* 1 *×* 10^5^; pink region shows worst to best scores) than the MCMC algorithm. The MCMC algorithm requires a great deal more time because it must perform hundreds of iterations to achieve a similar level of inference quality (mean score *∼ -*5 *×* 10^5^; standard deviation *∼* 8 *×* 10^5^; thick black line shows the mean, thinner black curves show 100 individual runs of the algorithm).

In summary, the EIG network which we describe below and in the main text reliably produces inferences that are as accurate as the best of these MCMC runs but far more quickly. EIG avoids the need for iterative computation by estimating 3D shape and texture properties via a single feedforward pass through a deep inference network. Further comparisons between MCMC and efficient inference networks for inverse graphics (using an earlier version of EIG, without the initial face detection stage and using a more limited training regime and loss function) can be found in^6^.

### EIG model

The EIG model is a multistage neural network that attempts to estimate the MAP (Maximum A Posteriori) 3D scene properties and identity of an observed face image (approximately maximizing the posterior in Equation 1 of the main text). EIG comprises three inference modules arranged in sequence to take advantage of the conditional independence structure in the generative (graphics) model. These three modules compute (1) a segmentation and normalization of the face image; (2) an estimate of the 3D face shape and texture; and (3) a classification of the individual whose face is observed.

Below we describe how each of these modules is constructed. The EIG network can also be seen as a multitask network that is designed to solve several tasks at once, including segmentation, 3D scene reconstruction, and identification, where the generative model determines which tasks should be solved and the conditional independence structure of the generative model determines the order in which they should be solved.

#### Estimating face image given a transformed image, Pr(*I|O*)

Given an observation consisting of a face image with cluttered background, *O*, MAP inference involves estimating *I∗* that maximizes Pr(*I|O*). This can be achieved by a segmentation of the observed image that only consists of the face-proper region and excludes the rest.

We implemented this inference problem using a convolutional neural network^7, 8^, referred to as *f*_1_ in the main text. We took a recent convolutional neural network with an hour-glass architecture that is trained for volumetric 3D segmentation of faces from images^9^. This model takes as input an image and outputs a 3D voxel map where a value of 1 indicates inside the face region and a value of 0 indicates outside the face region. The output of this network is a rough and noisy estimation of the face shape in the form of a voxel grid, *V*_*xyz*_, of dimensions 192 (width) *×*192 (height) *×*200 (depth), which we found in practice often includes filled but disconnected regions that are outside the face-proper region.

We adapted this output for accurate 2D segmentation of the face-proper region in the following way. We first sum over the depth dimension of *V*_*xyz*_ to obtain a 2D map, *V*_*xy*_, of dimensions 192 *×* 192. We then binarize *V*_*xy*_ (i.e., replace all non-zero entries with 1) and compute its connected components. We produce a segmentation of *O* using the largest connected region of *V*_*xy*_ as the mask. Finally, we normalize this region by zooming in on the segmented image using bicubic interpolation such that the resulting image’s longer dimension is 227. In practice, this procedure yields good estimates for *I∗*. We also applied a small amount of translation (25 pixels) away from left or right border for the normalized FIV images, which better aligned them with the samples from the generative model.

#### Scene parameters given face image, Pr(*S, T, L, P |I*)

Given a face image as input, MAP inference involves estimating the scene properties (latent variables in the graphics program), {*S∗, T ∗, L∗, P ∗*} maximizing Pr(*S, T, L, P |I*). We accomplish this using an inference model by learning to map inputs to their underlying latent variables in the graphics program.

Our inference model is a convolutional neural network with each layer implementing a cascade of functions including convolution, rectified linear activation, pooling, and normalization. We obtained this model by modifying Alexnet network architecture in the following way^10^: We removed its top two fully-connected layers and replaced them with a single new fully-connected layer. The details of the resulting network architecture is given in Extended Data Table 2.

We initialized the parameters of *f*_2_, *f*_3_, and *f*_4_ in the inference model using the corresponding weights of Alexnet that was pretrained on a large corpus of images, namely the Places dataset^11^. The pretrained network weights are provided by its authors and can be downloaded at http://places2.csail.mit.edu/models_places365/alexnet_places365.caffemodel. This dataset consists of about 2.5 million images and their corresponding place labels such as “beach,” “classroom,” “landscape,” etc. (365-way categorization). The parameters of the new fully-connected layer (also referred to as scene properties layer or latents layer) were initialized randomly. Using these pretrained weights ensured that the earlier layers of the inference model provided a good generic visual feature extractor not specifically related to faces. We also avoided using a face corpus pretrained weights as this would require access to a large labeled dataset of weights, which EIG doesn’t require.

To learn the mapping from images to their latent variable representations, we drew 200, 000 random samples from the generative model. Each resulting image was a 227 *×* 227 color image and each target was a concatenation of all the latent variables making a vector of length 404 (200 shape properties, 200 texture properties, and 4 extrinsic scene parameters). Half of the images were added background and were first segmented and normalized using *f*_1_, whereas the other half of the images were not added background and were directly used during training. We finetuned the parameters of *f*_3_, *f*_4_ starting from their pretrained weights and trained the parameters of *f*_5_ starting from random initialization. The network learns a mapping from these images to their latent variable representations, which we accomplish minimizing a mean squared error (MSE) loss function using stochastic gradient descent with minibatches of 20 examples. In our simulations we used a learning rate of 10^−4^. In order to ensure that gradients were large enough throughout training, we multiplied the target latent variable vectors by 10. We accounted for this preprocessing step by dividing the outputs of the network by 10 at test time. We trained the model for 75 epochs.

#### Person identity given scene parameters, Pr(*F |S, T, L, P*)

We provide the details of Pr(*F*) before describing this final component of the inference model. In principle, this distribution is over a finite set of familiar individuals but allowing for possibility of encountering a new, unfamiliar individual^12^. Here, we approximated Pr(*F*) as a uniform distribution over a set of familiar individuals. Specifically, we treated Pr(*F*) as a multinomial categorical distribution with *K* outcomes (i.e., *K* unique person identities) with each outcome equally probable. Each person identity is chosen as a pair of shape and texture properties and denoted as Pr(*S, T |F*).

Given scene properties, MAP inference involves estimating the person identity, *F ∗*, maximizing Pr(*F |S, T, L, P*). To estimate *F ∗* given scene properties, we extended the inference model with a new fully-connected layer, *f*_6_, of length *K*. To learn this mapping from scene properties to identities, we generated a new dataset of *K ∗ M* images where *M* is the number of times the shape and texture properties associated with each of the *K* identities were rendered. For each image, we randomly draw the lighting direction and pose properties from their respective prior distributions, Pr(*L*) and Pr(*P*). In our simulations we set *K* to 25 and *M* to 400.

For our FIV experiments, we do not have access to the ground truth shapes or textures of the 25 person identities and therefore we cannot use the graphics program for generating a training image set. Instead, for a given identity, we obtained *M* = 400 images by a bootstrapping procedure applied to the whole set of 7 attended face images for that identity. Given the image bounding box of the face proper region, we randomly and independently stretched or shrank each side of the bounding box by 15%. We resized the resulting bounding boxes by a randomly chosen scale between 75 - 99%. Finally, we translated the resulting bounding boxes in the image randomly but ensuring that the entire face-proper region remained in the image. We refer to the resulting image set as the bootstrapped FIV image set.

The training procedure was identical for the FIV and FIV-S experiments. We train the new identity classification layer *f*_6_ and finetune the scene properties layer *f*_5_ using *M ∗ K* = 10, 000 images and their underlying person identity labels minimizing cross-entropy loss^13^. We used a learning rate of 0.0005. We performed stochastic gradient descent with minibatches of 20 examples until the training performance was high (e.g., *>* 95%). In practice, it took two additional epochs of training for the FIV-S image set, and 20 additional epochs of training for the FIV image set.

All of our models are implemented using the pytorch machine learning library and will be made publicly available at the time of publication at https://github.com/iyildirim/efficient_inverse_graphics.

#### Weaknesses of EIG

We note two potential weaknesses of the inference model. First, it may not perform as well when the segmentation step *f*_1_ fails (e.g., too much of the background is left in the attended face image). We observed that this is an issue only if the face doesn’t cover a spatially significant portion of the input image. Second, the model’s reconstruction accuracy may degrade when the observed faces have shapes and textures far from the regions of high prior probability in the generative model, Pr(*S, T*). We see these weaknesses mostly as challenges for the model as currently implemented, with a rather limited set of face experiences for training compared to what an individual encounters over the course of their lifetime – let alone what is effectively a much broader base of experience over evolutionary time that also shapes the brain’s representations.

The training procedure underlying the third component of our inference model, Pr(*F |S, T, L, P*), helps alleviate the second issue by allowing finetuning of *f*_5_, thereby adjusting Pr(*S, T, L, P |I*) to the given training set (e.g., the bootstrapped FIV image set).

### VGG network

The VGG network is based on the raw pretrained VGG face network (referred to as VGG-Raw) that is publicly available, http://www.robots.ox.ac.uk/∼vgg/software/vgg_face/. This network consists of 13 convolutional layers (8 more layers than Alexnet) and 3 fully-connected layers (same as Alexnet). The dataset used for training this network consisted of more than 2.5 million face images where each image is labeled with one of 2622 person identities. The details of the network architecture, its training dataset and training procedure can be found in^14^.

Similar to the EIG network, the VGG network is obtained by finetuning this pretrained VGG-Raw network on the relevant image sets. For our FIV experiments, we used the same bootstrapped training dataset of FIV images as described above. We replaced VGG-Raw’s top 2622-way fully connected classification layer (i.e., its third fully-connected layer; TFCL) with a 25-way classification layer for the FIV identities. Training of VGG started from their pretrained values in VGG-Raw except this final layer which was initialized with random weights. We trained that new classification layer (TFCL) and finetuned the weights in TCL, FFCL, and SFCL using stochastic gradient descent to minimize a cross-entropy loss.

For our FIV-S experiments, we replaced the final classification layer in the pretrained VGG-Raw network with a 500-way classification layer. To train this network, we obtained a new dataset with the person identities and training images coming from the generative model. We first randomly sampled 500 identities as pairs of shapes and textures from *Pr*(*S, T |F*). We then rendered each identity using 400 viewing conditions randomly drawn from *Pr*(*L, P*), identical to EIG’s training dataset. This procedure gave us a total of 200, 000 images and their corresponding identity labels (from 1 to 500). In line with the training of the VGG-Raw network, the VGG network as well as the EIG network utilized two standard data augmentation methods including making an image grayscale with a low probability (0.1) and mirror reflecting an image with probability 0.5. As for our FIV experiments, we initialized the weights of the VGG network using the weights of the pretrained VGG-Raw network except for its classification layer, which was initialized using random weights. We then finetuned the weights associated with its TCL, FFCL, and SFCL and trained its classification layer using stochastic gradient descent to minimize a cross-entropy loss. We used a learning rate of 0.0001 with minibatches of size 20 images.

#### Evaluation of VGG-Raw, VGG^+^, and EIG^−^ networks on the FIV image set

The pretrained VGG-Raw network readily generalizes well to the FIV image set and gives rise to qualitative and quantitative patterns that are similar to the VGG network (compare Extended Data Fig. 2b-e to main text Fig. 2j-m).

In order to isolate contribution of EIG’s initial segmentation and normalization step (*f*_1_) to its better fit to neural data, we also tested VGG^+^ and EIG^−^ networks. The VGG^+^ network extends VGG with the initial segmentation and normalization step from the EIG network (*f*_1_; Extended Data Fig. 2f). Overall, the transformations across the layers of the VGG^+^ network (Extended Data Fig. 2g) were very similar to that of the VGG or VGG-Raw networks (Fig. 2j main text, Extended Data Fig. 2b). Even though VGG^+^ was still more view-invariant when compared to neural data (Extended Data Fig. 2h,i; *p <* 0.05 comparison of view-invariance coefficients for each layer), it was somewhat less view-invariant when compared to the VGG network (Fig. 2k,l main text) across all three of its layer indicating that the VGG network can pick up on spurious cues to discriminate identities (e.g., shirt color or hair) to some extent. Correlations between VGG^+^ and data similarity matrices were similar to the correlations based on the VGG network (compare Extended Data Fig. 2j and Fig. 2m main text) with the exception that the initial segmentation and normalization stage helped with VGG^+^’s fit to ML/MF: its TCL was more like ML/MF and less like AM in comparison to the VGG network’s TCL. These results suggest that even though the initial segmentation and normalization stage can help align the view-based TCL stage in the VGG network with the view-based ML/MF representations in the neural data, it is not sufficient to influence the later, more view-invariant stages of the network closer to the final classification step.

The EIG^−^ network is a lesion of the EIG network without the initial segmentation and normalization step. Even though this network replicates some of the qualitative structure seen in the EIG network, it does not capture the data nearly as well (compare Extended Data Figs. 2c and 2d), providing quantitative support for an initial attention-like step in the EIG network.

In Extended Data Fig. 3, we also present scatter plots showing the fine-grained relation between all pairwise similarities in each network layer and each neural site, for the EIG network (panel A) and the VGG network (panel B) in addition to showing how their earlier layers relate to neural data (panels C, D). See figure caption for discussion of these results.

### Neural data analysis

The neural experiments and the data presented in the main text were originally reported in^15^. Various relevant details can be found in SI Section 2.

#### Representational similarity matrices: Neurons

To compute the neural similarity matrices for a given neural site, each image was represented as a vector of the average spiking rates of all neurons recorded at that site. Following^16^, we obtained the average number of spikes for each neuron across the repetitions of a given image using the time-binned spike counts centered at 200 msec after stimulus onset with a time window of 50 msec in each direction. Following^15^, for each site, we min-max (range [0, 1]) normalized the average spiking rate of each neuron. For a given neural site, similarity of a pair of images was computed as the Pearson’s correlation coefficient of the corresponding pair of the average spiking vectors. All spiking data was processed using the Neural Decoding Toolbox^17^.

#### Representational similarity matrices: Models

For a given image set, model, and the model’s layer, images were represented as a vector of activations of all units in that layer. The model similarity of a pair of images (e.g., each entry in the similarity matrix shown in Fig. 2f, main text) is the Pearson’s correlation coefficient of their corresponding activations vectors.

#### Linear regression analysis using the idealized similarity templates

For a given representational similarity matrix *M*, we solved the following linear equation.

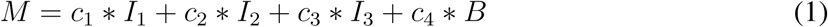

where {*c*_1_, *c*_2_, *c*_3_, *c*_4_} are coefficients, *I*_1_ is the idealized view-specificity matrix, *I*_2_ is the idealized mirror-symmetry matrix, *I*_3_ is the idealized view-invariant identity coding matrix, and *B* is the background matrix. These matrices are shown in Fig. 2e in the main text. All black entries have a value of 1, all gray entries have a value of 0.5 and all white entries have a value of 0. We solve this equation using a non-negative least squares solver as implemented in the Python package `scipy`’s `nnls` method.

#### Bootstrap procedure

Due to the small number of subjects (N=3), we performed bootstrap analysis at the image-level. Following the procedure in^18^, a bootstrap sample was obtained by sampling the 175 images in the FIV image set with replacement. Based on this sample, we computed the neural and the model similarity matrices. To avoid spurious positive correlations, we excluded all non-diagonal identity-pairs that could arise due to sampling-with-replacement. Based on the discussion in^19^ and following^20^, we computed the Pearson correlation coefficient between pairs of representational similarity matrices. We repeated this procedure for 10, 000 bootstrap samples. Significance was measured using a direct bootstrap hypothesis testing procedure with a significance level of 0.05.

For the linear regression analysis with idealized similarity matrices, we again bootstrap sampled the 175 images with replacement and performed the linear regression using the resulting similarity matrix. We repeated this procedure for 10000 times.

### Psychophysics methods

#### Experiment 1

The experimental procedure consisted of a simple “same”/”different” judgment task as the following. A study item was presented for 150 msecs, which was followed by a masking stimuli in the form of a scrambled image of a face for 500 msecs. Finally a test item appeared and stayed on until a response was entered (the participants were instructed to press “f” for “same” and press “j” for “different”). They performed 10 practice trials before performing 96 experimental trials. Participants did not receive any feedback at all during the practice trial, which aimed to have participants get used to the experiment parameters (e.g., its interface). During the experimental trials, participants were shown their current average performance at every fifth trial.

The stimuli were 200 *×* 200 color images of faces photo-realistically rendered using the generative model. None of the stimuli across the experiments were used during training of the models. The viewing conditions for both the study and test items were drawn randomly from their respective prior distribution, *Pr*(*L, P*). All participants saw the same image set (i.e., the viewing conditions were sampled once for all participants before the experiment began). There were 48 “same” trials and 48 “different” trials.

No study identity (i.e., a pair of shape and texture properties) was presented twice across trials. For the “different” trials, we chose the distractor face (the test item) by running a Metropolis-Hasting based search until 50 accepted steps. The search started from a random face but with matching lighting and pose parameters as that of the study item and increasingly moved closer to the study face w.r.t. likelihood *P* (*I|S, T, L, P*) by generating proposals from the prior distribution over shape and texture properties, *Pr*(*S, T*). This procedure aimed to ensure that the test facial identities in “different” trials were not arbitrarily different from the study item in obvious ways. Our data suggested that this procedure was effective: across the “different” trials average Pr(“Same”) was 0.35 with a standard deviation of 0.15, min value of 0.10 and max value of 0.71. All stimuli were rendered using Matlab’s OpenGL-based rendering pipeline.

#### Experiment 2

The stimuli and procedure were identical to Experiment 1 with the following exceptions. The test item was always presented frontal (i.e., frontal lighting and frontal pose) and without texture. This was achieved by assuming a uniform gray color for all vertices of the face mesh before rendering.

#### Experiment 3

The stimuli and procedure were identical to Experiment 1 with the following exception. The test item was always presented frontal (i.e., frontal lighting and frontal pose), however, the texture was rendered on a flat surface in order to eliminate shape information from shading. In an attempt to further eliminate the shape information, we post-processed the resulting images by applying a fish-eye lens effect. All of the code used to generate stimuli as well as the actual images used in the experiments will be released at the time of publication.

#### Calculating Similarity(study,test)

For a given pair of study and test images, their predicted similarity by a model was computed as the similarity of their respective representations under the model. For the EIG network, this representation was its *f*_5_ consisting of the shape and texture properties (a 400 dimensional vector), excluding the lighting and pose parameters. The model’s similarity prediction was the Pearson’s correlation coefficient of these two vectors.

For the other networks, the images were represented by their resulting SFCL activations. The model’s prediction is the correlation coefficient of these two vectors. We found that no other layer in the VGG network resulted in a better account of the human behavior than the layer we used. We also considered using other similarity metrics in addition to Pearson’s correlation coefficient such as the cosine of the angle between two vectors and Euclidean distance. We found no significant difference in fits for any of the models.

#### Lighting elevation judgment task

Both groups of participants had to complete 5 training trials before they moved onto 45 test trials. We only used the test trials in our analysis. Each of the 45 trials featured a different facial identity. In the depth-suppression group, each of the 9 levels of depth suppression (from 1, regular faces, to 0, flat face, to −1, fully inverted faces with nose pointing away from the observer; see also the main text) appeared 5 times throughout the experiment. In the lighting source elevation experiment, each of the 9 levels of elevation appeared 5 times (from the top of the face, 1.31 radians of elevation, to the front of the face, 0 radians of elevation, to the bottom of the face, −1.31 radians of elevation; see also the main text).

For each condition, we z-scored each participant’s responses (a total of 45 ratings each in the range of 1 to 7) before averaging all responses across participants and across the 9 levels. The error bars were obtained for each of the 9 levels as the standard deviation of the average values of the 5 stimuli items corresponding to that level.

Obtaining the EIG network’s predictions was straightforward. For each condition, we ran the EIG model on the same set of 45 images as the human subjects, recording its outputs for the lighting elevation, *L*_*e*_. We averaged the values for the 5 images of each of the 9 levels. The error bars in the main text (Fig. 5c) show the standard deviation across these five images. The main text also reports trial-level correspondence between the model and the behavior as the correlation of model’s predicted lighting elevations and the average human response per each of the 90 test trials (Fig. 5d).

#### Face depth judgment task

Before the beginning of the experimental trials, participants were instructed that they would see frontal images of faces and some faces would be more flatter than others. They were shown several examples of fairly flat and fairly deep faces, which were samples chosen from either tail of the flatness distribution of 3000 randomly generated heads. On a given trial, participants were presented frontal image of a face (excluding neck and the ear) and were asked to judge the profile depth of it using a scale of 1 to 7. Next to the flat-end (wide-end) of the continuum participants were presented with the profile view of an altered mean-MFM-face with its depth scaled to −3 (+3) standard deviations away from the mean depth of the above mentioned 3000 faces. An example trial in this experiment is shown in Extended Data Fig. 12.

Participants had to complete 10 training trials before they moved onto 108 test trials. We only used the test trials in our analysis. The 108 trials featured 54 different facial identities with each identity rendered once as a regular face and once with depth suppression. These identities were uniformly assigned to the 9 depth suppression levels (6 identities per level). When rendering an identity as a regular face, we set the lighting elevation location to match where it would be perceived given its depth-suppression level according to the results in Fig. 5c in main text. The actual values used are indicated in the x-axis of Fig. 5e right panel. When rendering an identity with depth suppression we always place the lighting elevation at top, at 1.31 radians. Following previous work^21^, we rendered only the face proper region excluding ears and neck. This procedure resulted in 6 images per each of the 9 depth-suppression levels and 6 images per each of the 9 control levels.

For each condition, we z-scored each participant’s responses (a total of 108 ratings each in the range of 1 to 7) before averaging all responses across participants. The error bars were obtained for each of the 9 levels as the standard deviation of the average values of the 5 stimuli items corresponding to that level.

The EIG network can be readily used to estimate depth of a given face image. We ran the EIG model on the same set of 108 images as the human subjects, recording its outputs for the shape parameters, *S*. We then assigned a depth for each input image as the average displacement of the three key points (nose, left cheek, and right cheek) of the face shape with respect to the underlying aligned coordinate system of MFM. This coordinate system is in arbitrary units so we z-scored model’s predictions to bring it to the same scale as the behavioral data. The error bars in the main text (Fig. 5e, main text) show the standard deviation across six images falling under the same pair of depth-suppressed or control and one of the 9 levels. The main text also reports trial-level correspondence between the model and the behavior as the correlation of model’s predicted depth and the average human response per each of the 108 test trials (Fig. 5f, main text).

## Acknowledgements

This work was supported by the Center for Brains, Minds and Machines (CBMM), funded by NSF STC award CCF-1231216; the National Eye Institute of NIH (R01 EY021594 to W.A.F.); The New York Stem Cell Foundation (to W.A.F.); ONR MURI N00014-13-1-0333 (to J.B.T.); a grant from Toyota Research Institute (to J.B.T.); and a grant from Mitsubishi MELCO (to J.B.T.). W.A.F. is a New York Stem Cell FoundationRobertson Investigator. A high performance clustering environment for computations (Openmind) was provided by the McGovern Institute for Brain Research. The content is solely the responsibility of the authors and does not necessarily represent the official views of NIH.

## Supplementary Information

Supplementary Information can be found at the end of this document.

**Extended Data Table 1:** Pose distributions for the FIV-S image set (in radians).

| Pose Category | Azimuth ($P_z$) | Elevation ($P_x$) |
| --- | --- | --- |
| Frontal | $N(0, 0.05)$ | $N(0, 0.05)$ |
| Right-half profile | $0.75 + N(0, 0.05)$ | $N(0, 0.05)$ |
| Right profile | $1.50 + -1 * abs(N(0, 0.05))$ | $N(0, 0.05)$ |
| Left-half profile | $-0.75 + N(0, 0.05)$ | $N(0, 0.05)$ |
| Left profile | $-1.50 + abs(N(0, 0.05))$ | $N(0, 0.05)$ |
| Up | $N(0, 0.05)$ | $0.5 + N(0, 0.05)$ |
| Down | $N(0, 0.05)$ | $-0.5 + N(0, 0.05)$ |

**Extended Data Table 2:**
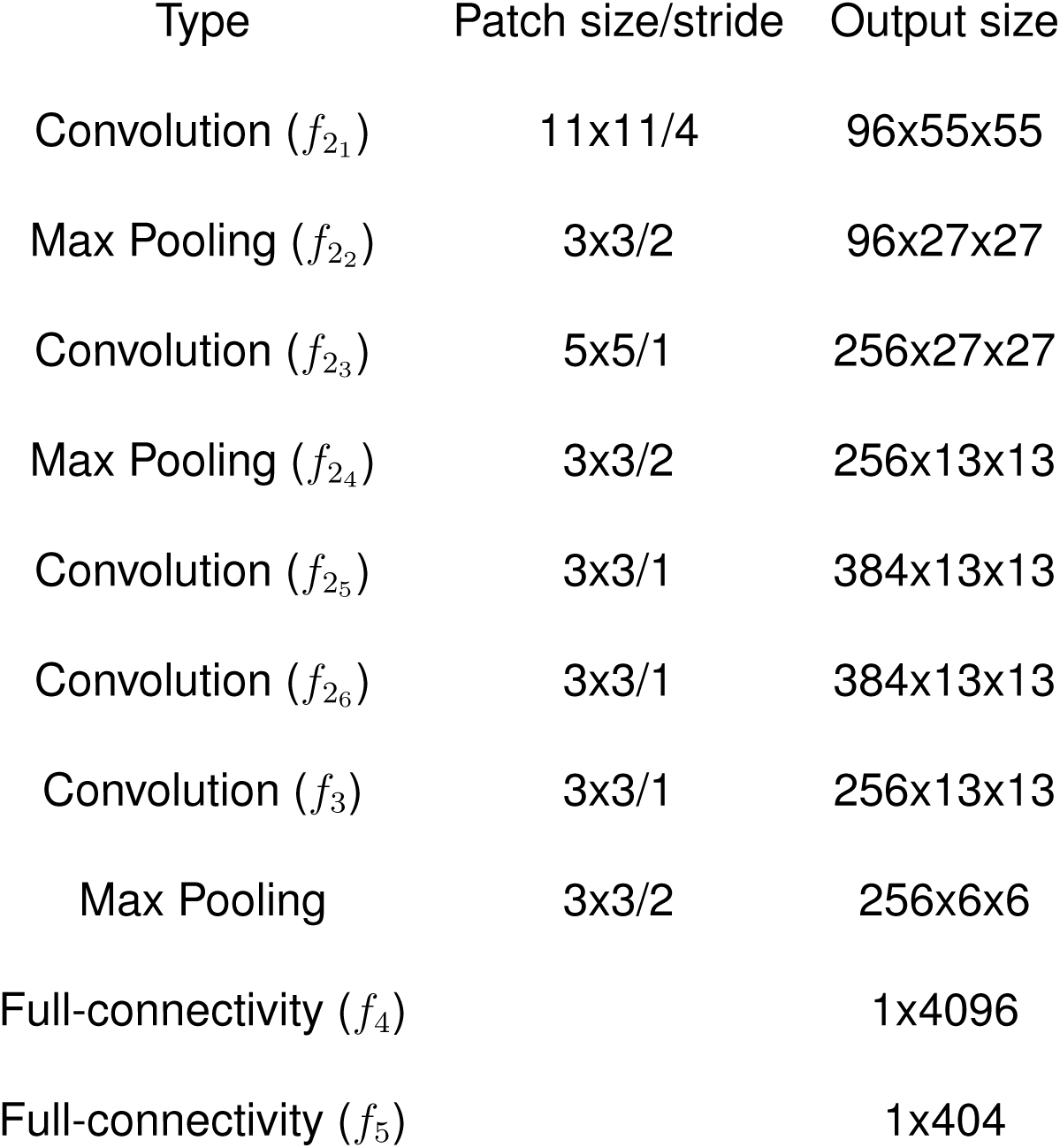
Inference model architecture

**Extended Data Table 3:** ID network architecture

| Type | Patch size/stride | Output size |
| --- | --- | --- |
| Convolution | 11x11/4 | 96x55x55 |
| Max Pooling | 3x3/2 | 96x27x27 |
| Convolution | 5x5/1 | 256x27x27 |
| Max Pooling | 3x3/2 | 256x13x13 |
| Convolution | 3x3/1 | 384x13x13 |
| Convolution | 3x3/1 | 384x13x13 |
| Convolution (TCL) | 3x3/1 | 256x13x13 |
| Max Pooling | 3x3/2 | 256x6x6 |
| Full-connectivity (FFCL) |  | 1x4096 |
| Full-connectivity (SFCL) |  | 1x4096 |
| Full-connectivity (TFCL) |  | 1x500 |

**Extended Data Table 4:** VAE decoder architecture

| Type | Patch size/stride | Output size |
| --- | --- | --- |
| 2D De-convolution | 4x4/1 | (64*8)x4x4 |
| 2D De-convolution | 4x4/2 | (64*4)x8x8 |
| 2D Batch-Norm |  | (64*4)x8x8 |
| 2D De-convolution | 4x4/2 | (64*2)x16x16 |
| 2D Batch-Norm |  | (64*2)x16x16 |
| 2D De-convolution | 4x4/2 | (64*1)x32x32 |
| 2D Batch-Norm |  | (64*1)x32x32 |
| 2D De-convolution | 4x4/2 | 3x64x64 |
| Sigmoid |  | 3x64x64 |

**Extended Data Table 5:** VAE-QN Pose architecture

| Type | Patch size/stride | Output size |
| --- | --- | --- |
| Convolution | 3x3/2 | 256x13x13 |
| Convolution | 3x3/2 | 256x7x7 |
| Convolution | 2x2/1 | 256x1x1 |
| Full-connectivity |  | 1x1x2 |
| TanH |  | 1x1x2 |

**Extended Data Fig. 1:**
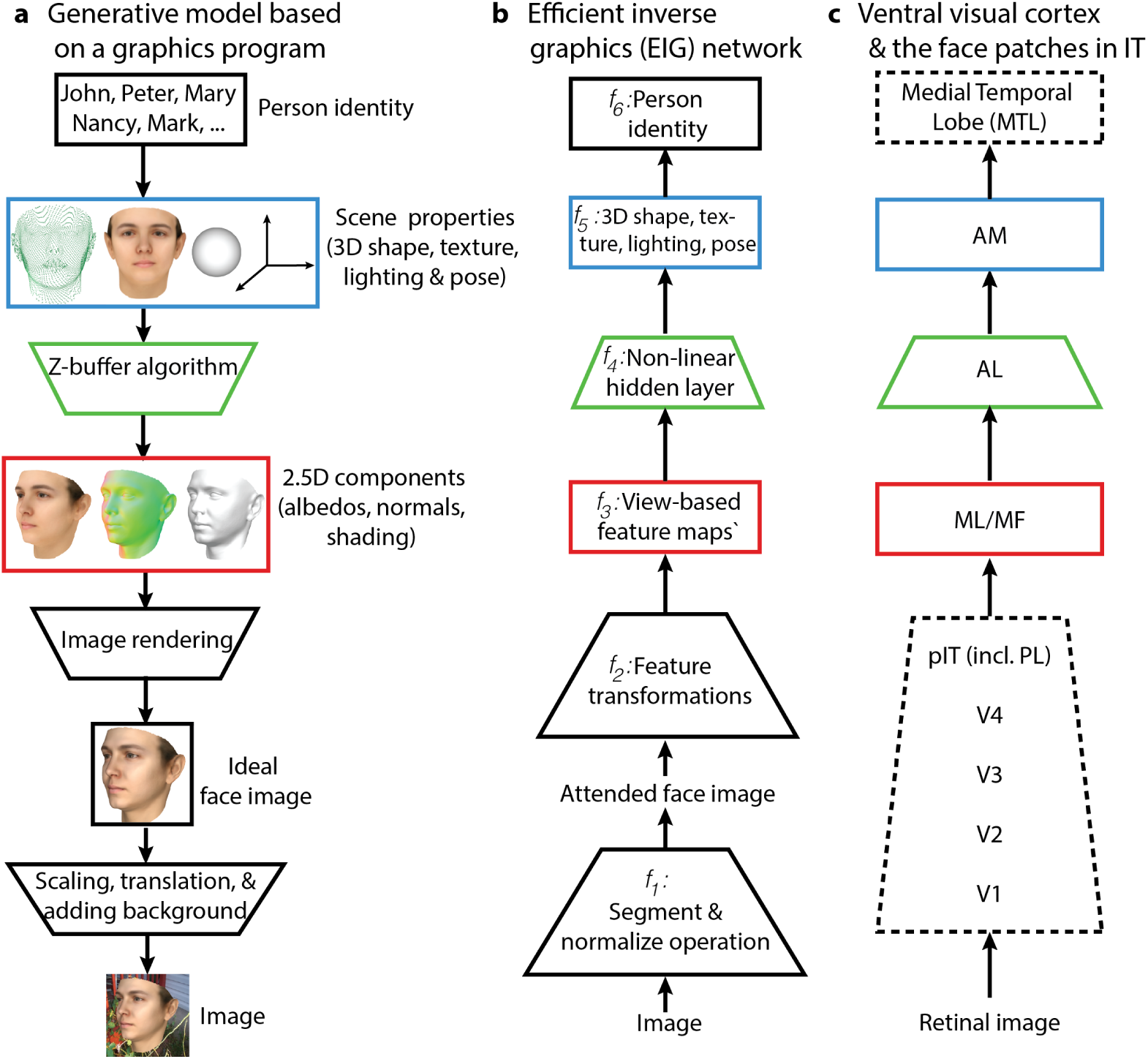
A more detailed diagram of the modeling framework. **a**, Schematic of the probabilistic generative that consists of a distribution over familiar person identities, detailed view of a graphics program with its key stages exposed, and an image-level transformations module. **b**, Schematic of the efficient inverse graphics model for efficient inference in the generative model based on DCNNs. *f*_1_ is for face segmentation and normalization, *f*_2_ to *f*_5_ for 3D scene parameter inference, and finally *f*_6_ for person identity recognition. **c**, Schematic of the ventral stream hierarchy with the three face patches indicated. Colored boxes in (**a**) to (**c**) show the hypothesized explanations of the neural sites based on the generative and inference models. Rectangles indicate representations, trapezoids indicate transformations or algorithms using representations.

**Extended Data Fig. 2:**
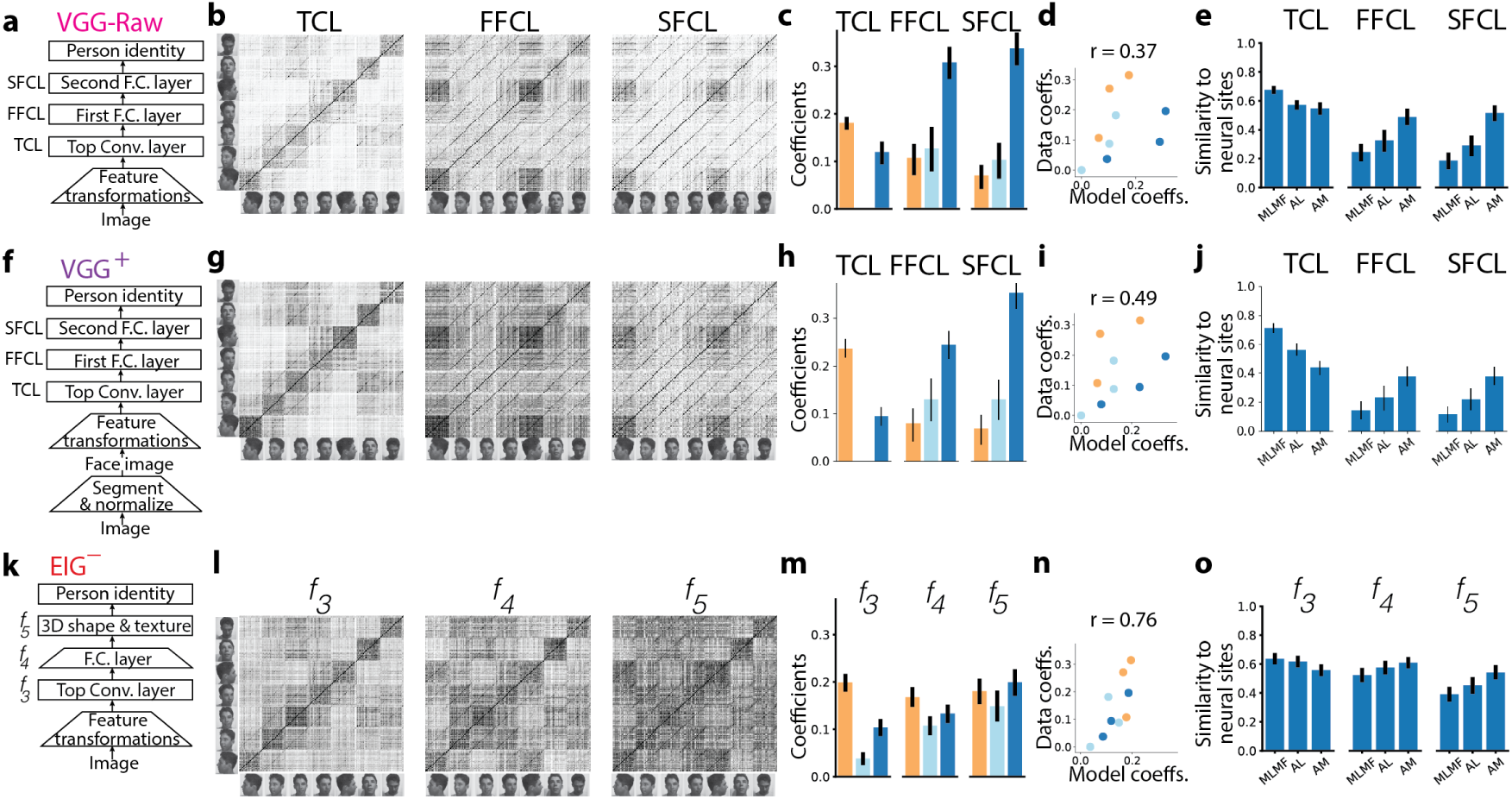
Evaluation of VGG-Raw, VGG^+^, and EIG^−^ networks based on the FIV image set (Extending Fig. 2 from main text). **a**, A schematic of the VGG-Raw network architecture. **g**-**j**, Analysis of the VGG-Raw network. **f**, A schematic of the VGG^+^ network architecture. Unlike (a), this network has an initial segmentation and normalization step. **g**-**j**, Analysis of the VGG^+^ network. **k**, The EIG^−^ network architecture. **l**-**o**, Analysis of the EIG^−^ network. Each row of the figure follow the same convention consisting of five parts (from left to right): (i) network architecture, (ii) similarity matrices of the key model layers, (iii) linear coefficients based on idealized similarity template analysis, (iv) comparison of data and model coefficients, and (v) comparison of data and model similarity matrices. Error bars show 95% bootstrap confidence intervals (CIs).

**Extended Data Fig. 3:**
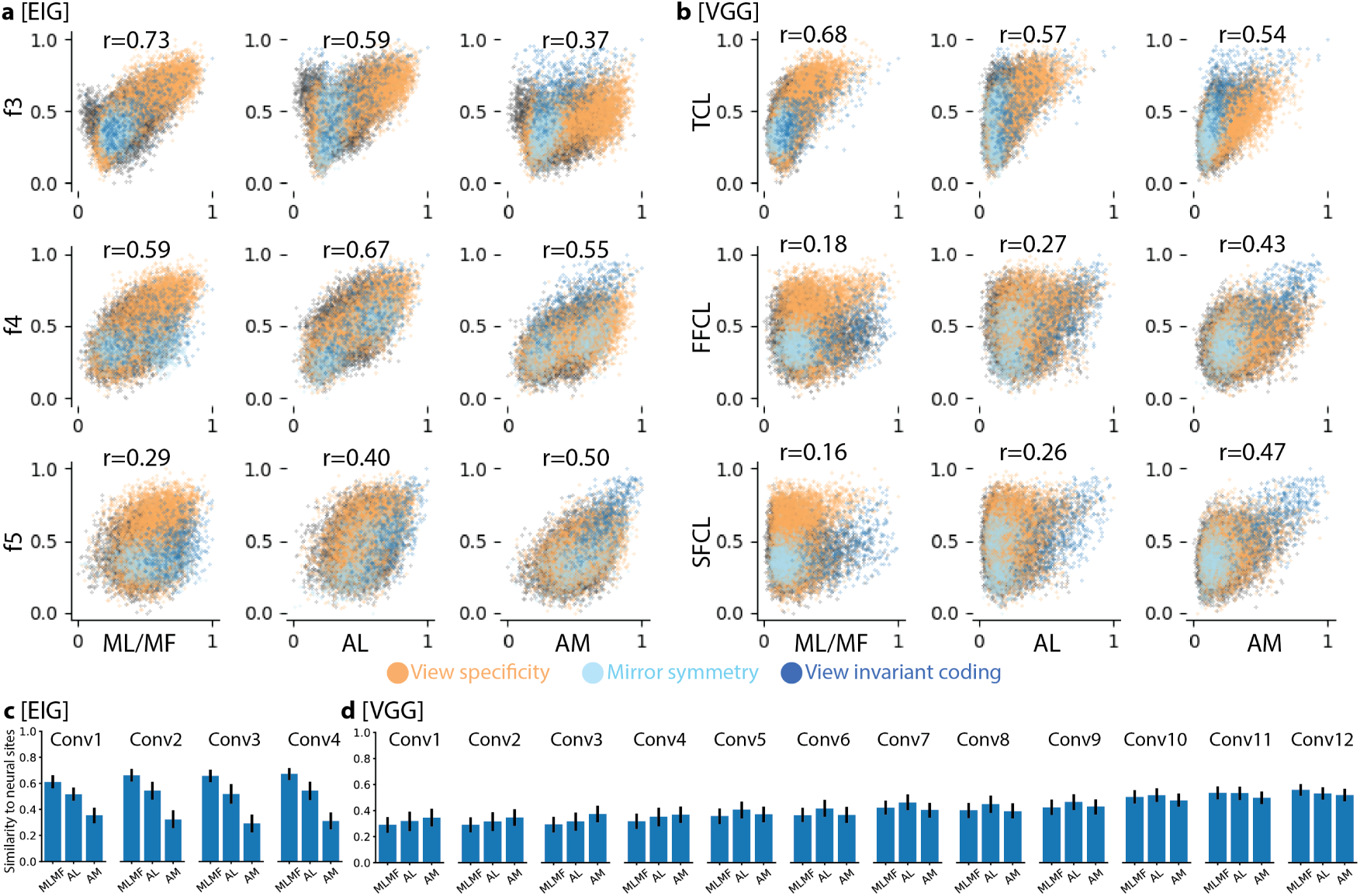
Scatter plots of data and model similarity matrices and analysis of earlier network layers (Extending Fig. 2 from main text). **a**, Scatter plots for every pair of EIG model layer and neural site. **b**, Scatter plots for every pair of VGG network layer and neural site. In (**a**) and (**b**), individual dots are colored according to whether they represent pairs of images showing the same view of different faces (yellow), the same face in different views (dark blue), or mirror-symmetric views of different faces (light blue). We can see that the quantitatively presented results in Fig. 2i and 2m are qualitatively visible in these scatter plots. For example, EIG’s layer *f*_3_ shows the strongest linear relationship with ML/MF, *f*_4_ shows the strongest linear relationship with AL, and *f*_5_ shows the strongest linear relationship with AM. Moreover, color-coding shows how the network layers which best capture the qualitative coding preferences in each face patch also best capture fine-grained structure within those qualitative categories: EIG *f*_3_ best captures the fine-grained structure of view-dependent coding in ML/MF, EIG *f*_4_ best captures the finegrained structure among mirror-symmetric views in AL, and EIG *f*_5_ best captures the fine-grained structure of view-invariant identity encoding in AM. **c**, Pearson’s r between the similarity matrices arising from the earlier layers in the second stage of the EIG network and neural sites. **d**, Pearson’s r between the similarity matrices arising from the earlier layers in the VGG network and neural sites. In (**c**) and (**d**), we see that the basic differences observed between EIG and VGG in their top convolutional layers are also seen at earlier convolutional layers, and none of the earlier layers in either network provide better accounts of ML/MF than the top convolutional layers. Error bars show 95% bootstrap confidence intervals (CIs).

**Extended Data Fig. 4:**
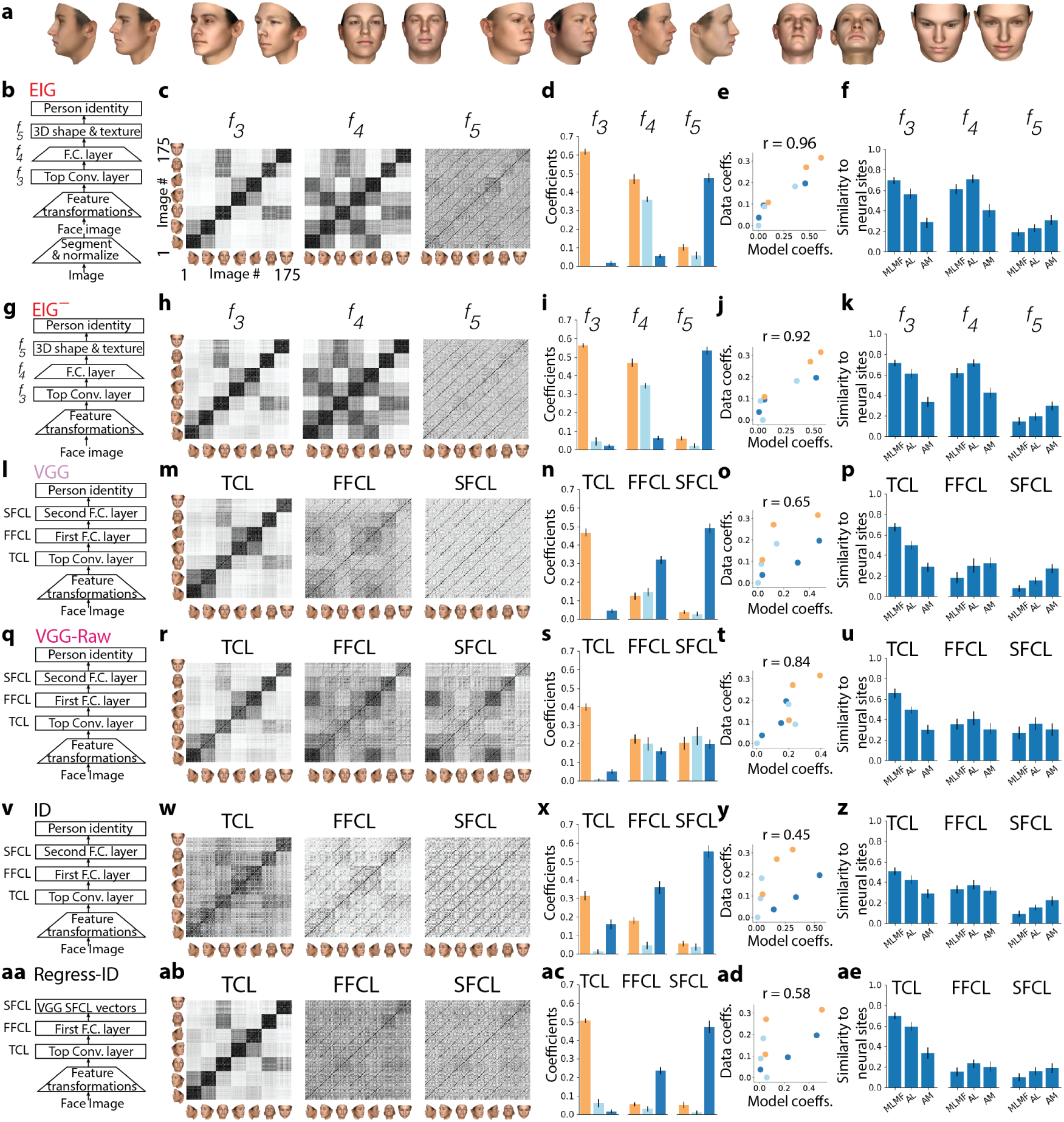
Evaluation of alternative models using the FIV-S image set. **a**, Sample images from the FIV-S image set. **b**-**f**, Results of the full EIG network. **g**-**k**, Results of the EIG^−^ network. **l**-**p**, Results of the VGG network. **q**-**u**, Results of the VGG-Raw network. **v**-**z**, Results of the ID network. **aa-ae**, Results of the Regress-ID network. Each row of the figure follows the same convention as in the Extended Data Fig. 2. Error bars show 95% bootstrap confidence intervals (CIs).

**Extended Data Fig. 5:**
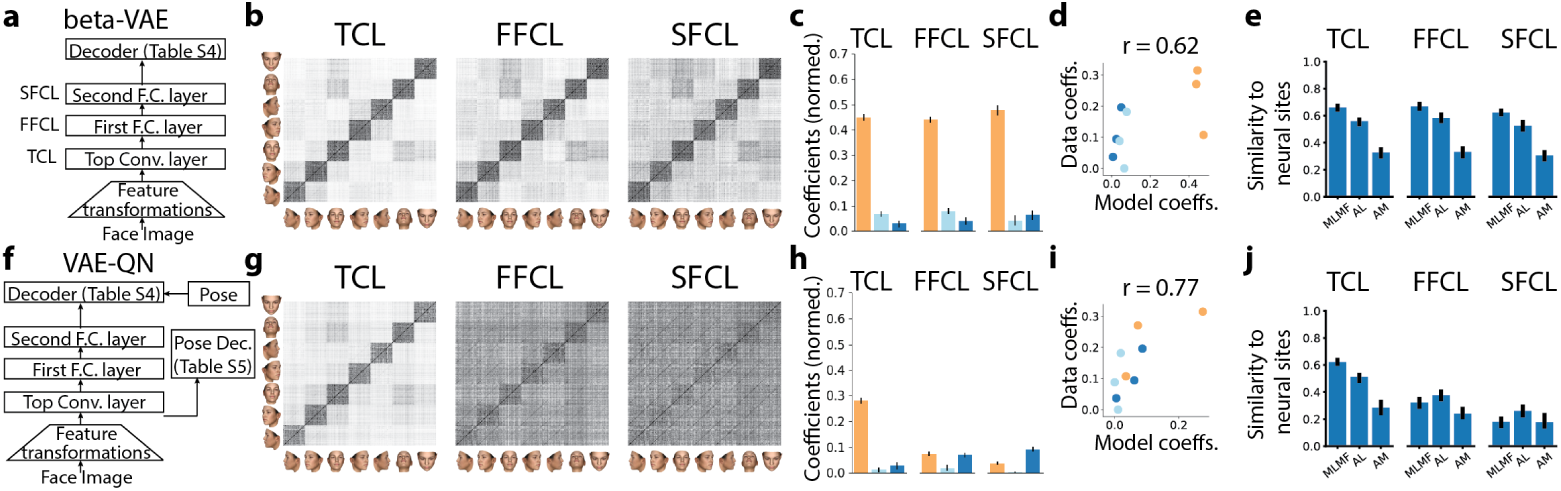
Evaluation of the VAE models using the FIV-S image set. **a**-**e**, Results of the betaVAE model. **f**-**j** Results of the VAE-QN network. Each row of the figure follows the same convention as in the Extended Data Fig. 2. Error bars show 95% bootstrap confidence intervals (CIs).

**Extended Data Fig. 6:**
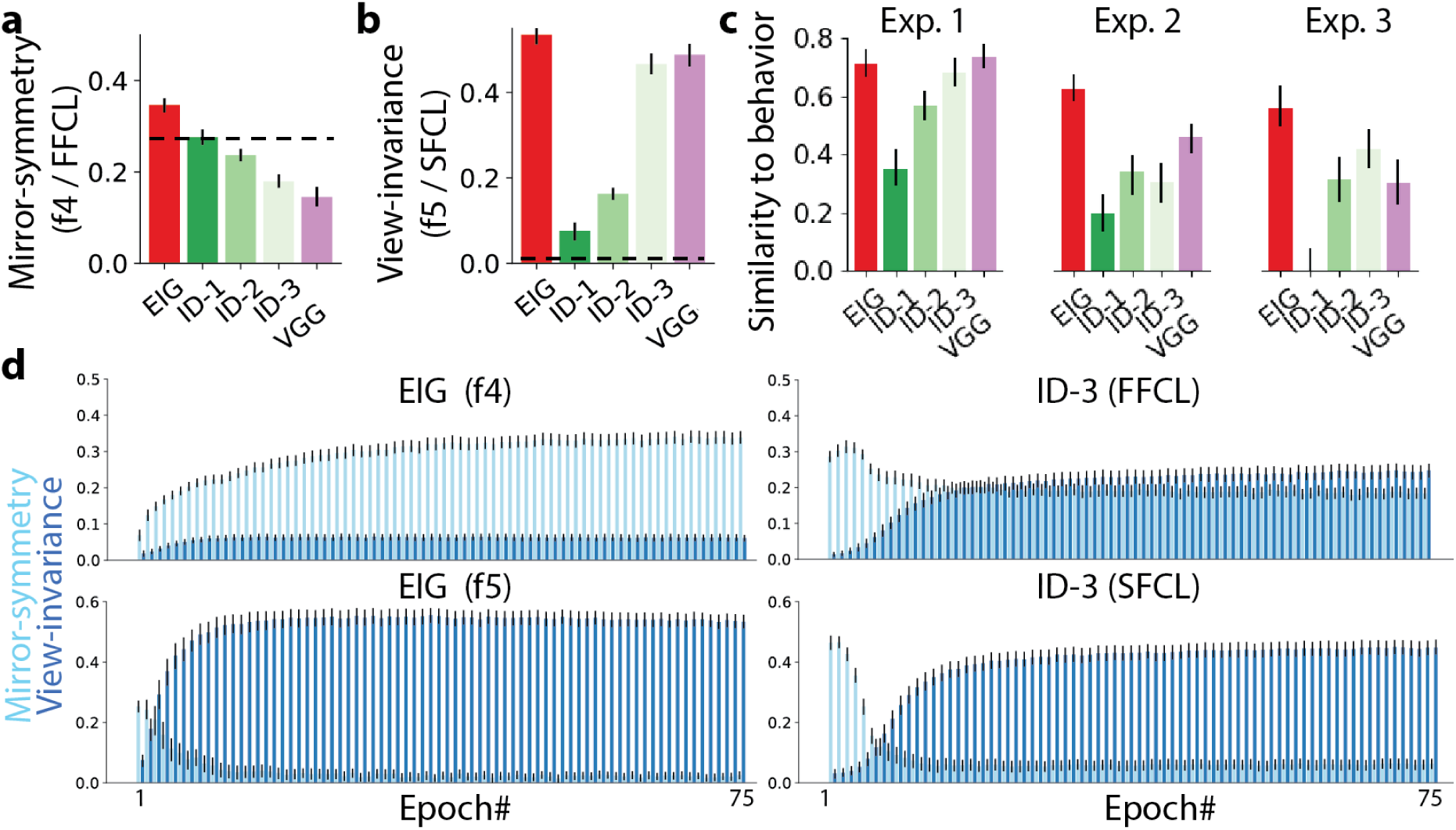
Trade-off arising from training targets and using pretrained weights. **a**, Mirror-symmetry coefficients in *f*_4_ of the EIG network, FFCL of the three variants of the ID network (ID-1, ID-2, ID-3), and FFCL of the VGG network. **b**, View-invariance coefficients in *f*_5_ of the EIG network, SFCL of the three variants of the ID network, and SFCL of the VGG network. **c**, Similarity to behavior for each behavioral experiment and model pair (extending Fig. 4b, main text). **d** Evolution of mirror-symmetry and view-invariance coefficients throughout training in the EIG network’s *f*_4_ and *f*_5_ layers (left column) and the ID-3 network’s FFCL and SFCL layers (right column). Error bars show 95% bootstrap confidence intervals (CIs).

**Extended Data Fig. 7:**
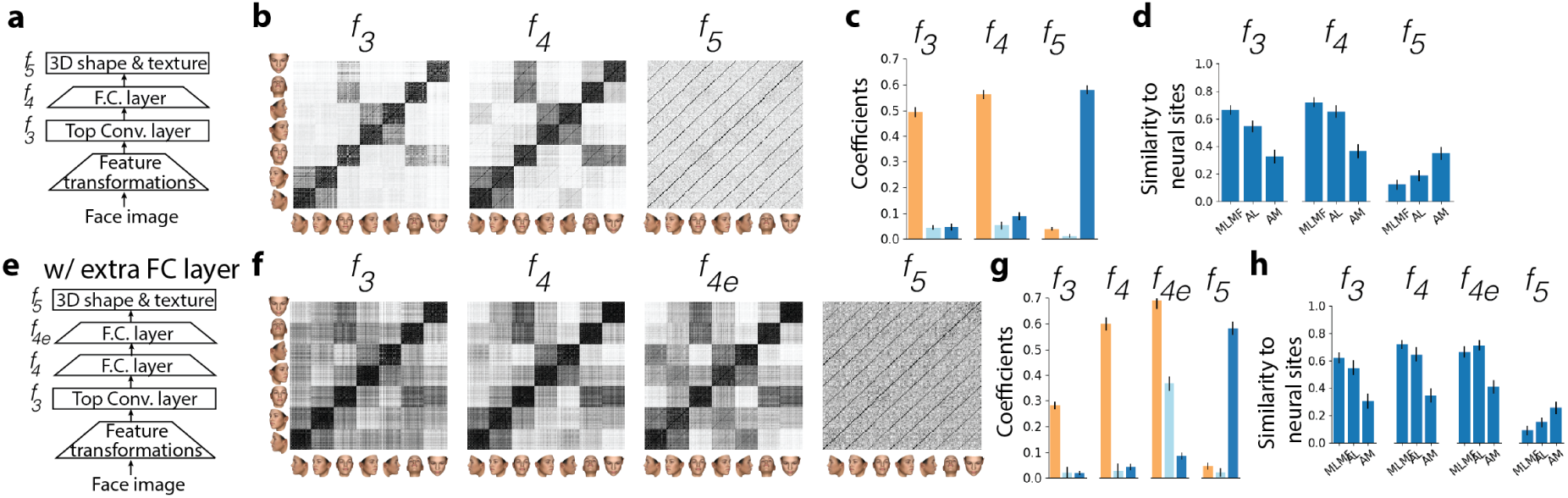
Variants of the EIG network architecture each trained from scratch without pretraining. **a**-**d**, The regular EIG architecture with a single hidden full-connectivity layer tested using the FIV-S image set. Notice that mirror-symmetry in the intermediate stage is significantly reduced. **e**-**h**, The EIG architecture extended with an extra hidden full-connectivity layer (*f*_4*e*_) and tested using the FIV-S image set. With this additional stage (which equates the total number of hidden layers in this network to that found in the ID network), this network gives rise to mirror-symmetry even though it is trained from scratch. Its second hidden full-connectivity layer (*f*_4*e*_; **f**, **g**) shows strong mirror-symmetry. Each row of the figure follows the same convention as in the Extended Data Fig. 2. Error bars show 95% bootstrap confidence intervals (CIs).

**Extended Data Fig. 8:**
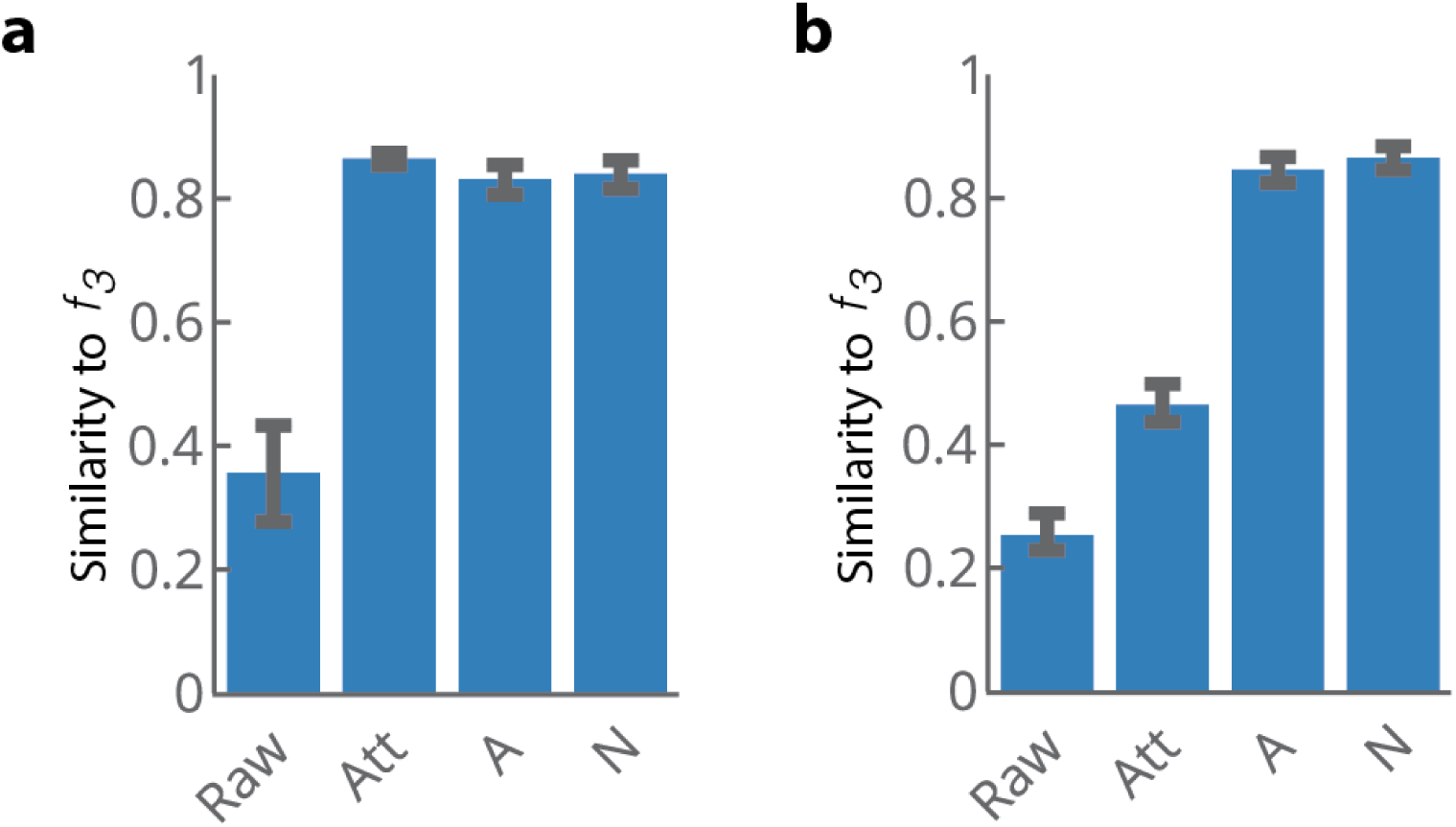
Comparison of intermediate stages of the generative model to *f*_3_. **a**, FIV image set based comparisons of *f*_3_ similarity patterns to that of raw images, attended images, and the 2.5D components. **b**, FIV-S-2 image set based comparisons of *f*_3_ similarity patterns to that of raw images, attended images, and the 2.5D components.

**Extended Data Fig. 9:**
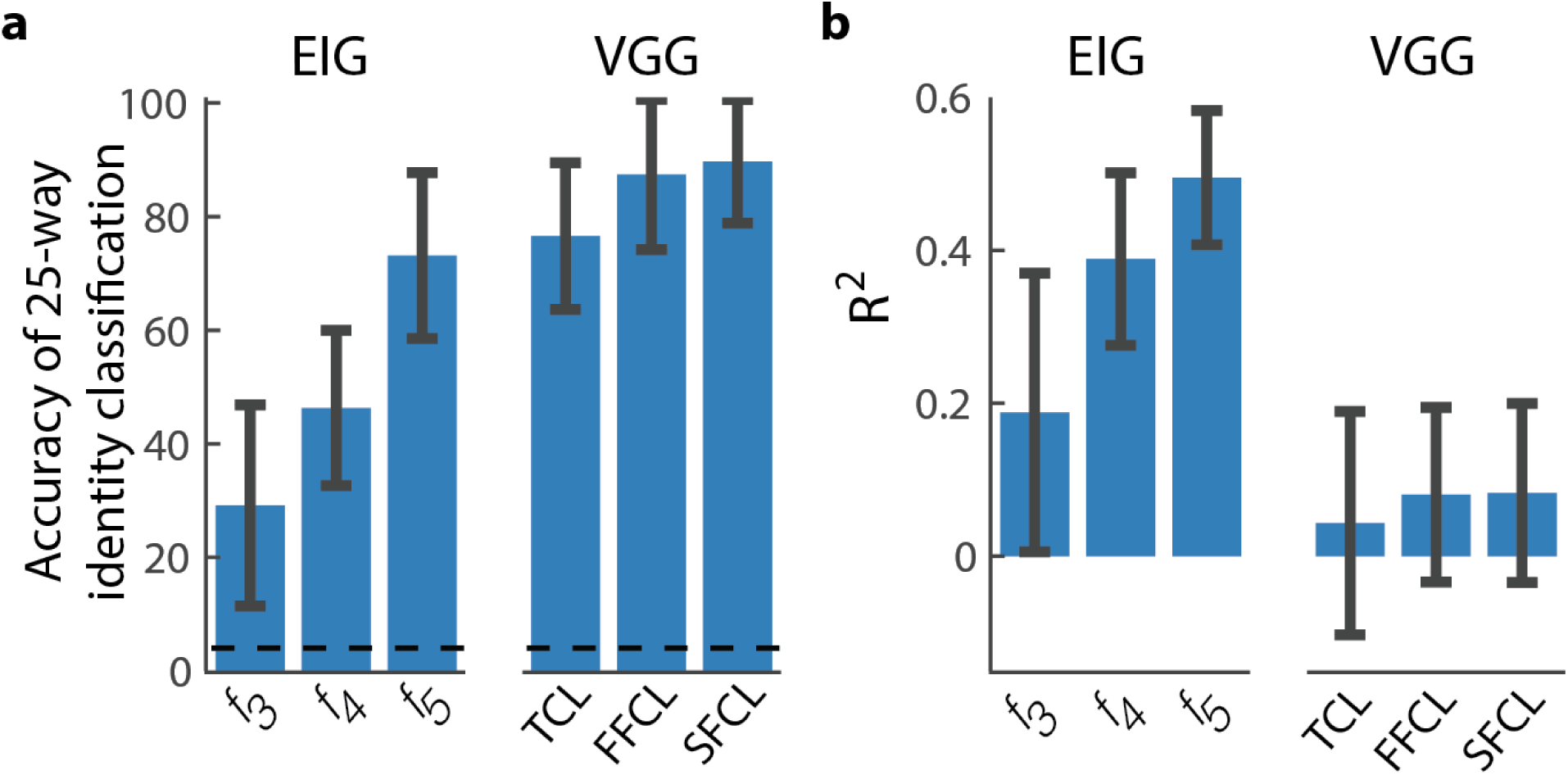
Decoding analysis. **a**, Average accuracy of a 25-way linear classifier decoding FIV identities from the VGG-Raw network and the EIG network. Dashed line shows chance performance (4%). **b**, Average goodness-of-fit *R*^2^ values resulting from linearly decoding approximate shape and texture properties of the FIV images from the VGG-Raw network and the EIG network. Error bars indicate standard deviation. All results are based on held-out test sets (see text for further details).

**Extended Data Fig. 10:**
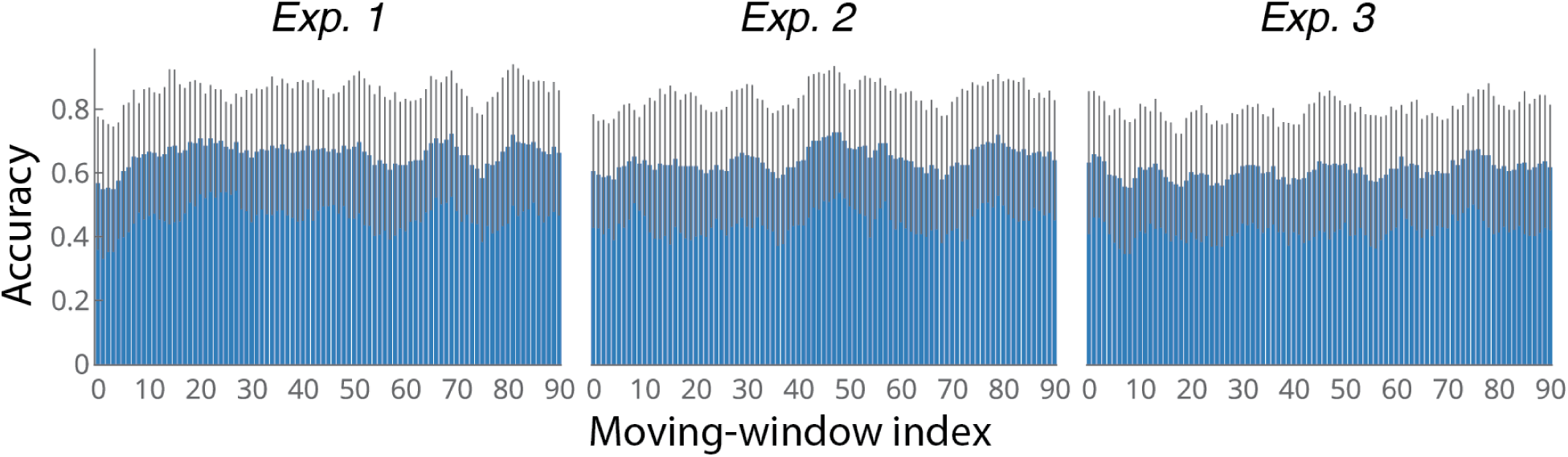
Learning curve analysis. The moving-window average performance of the participants in each experiment. We don’t observe any pronounced effects of learning, especially in Exps. 2 and 3. Error bars indicate one standard deviation.

**Extended Data Fig. 11:**
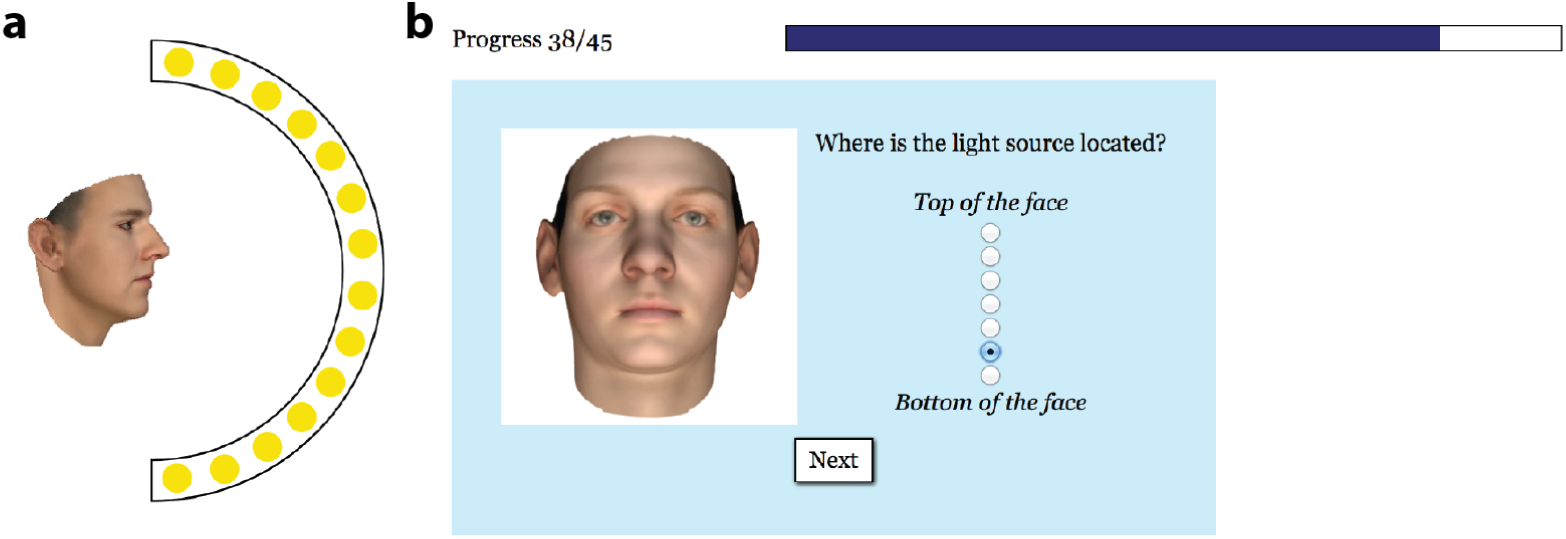
Lighting direction judgment experiment. **a**, The lighting source could be located at one of the 9 locations frontal to the center of the face, as illustrated they covered the full range from above the face (1.31 rads) to below the face (−1.31 rads). **b**, An example trial.

**Extended Data Fig. 12:**
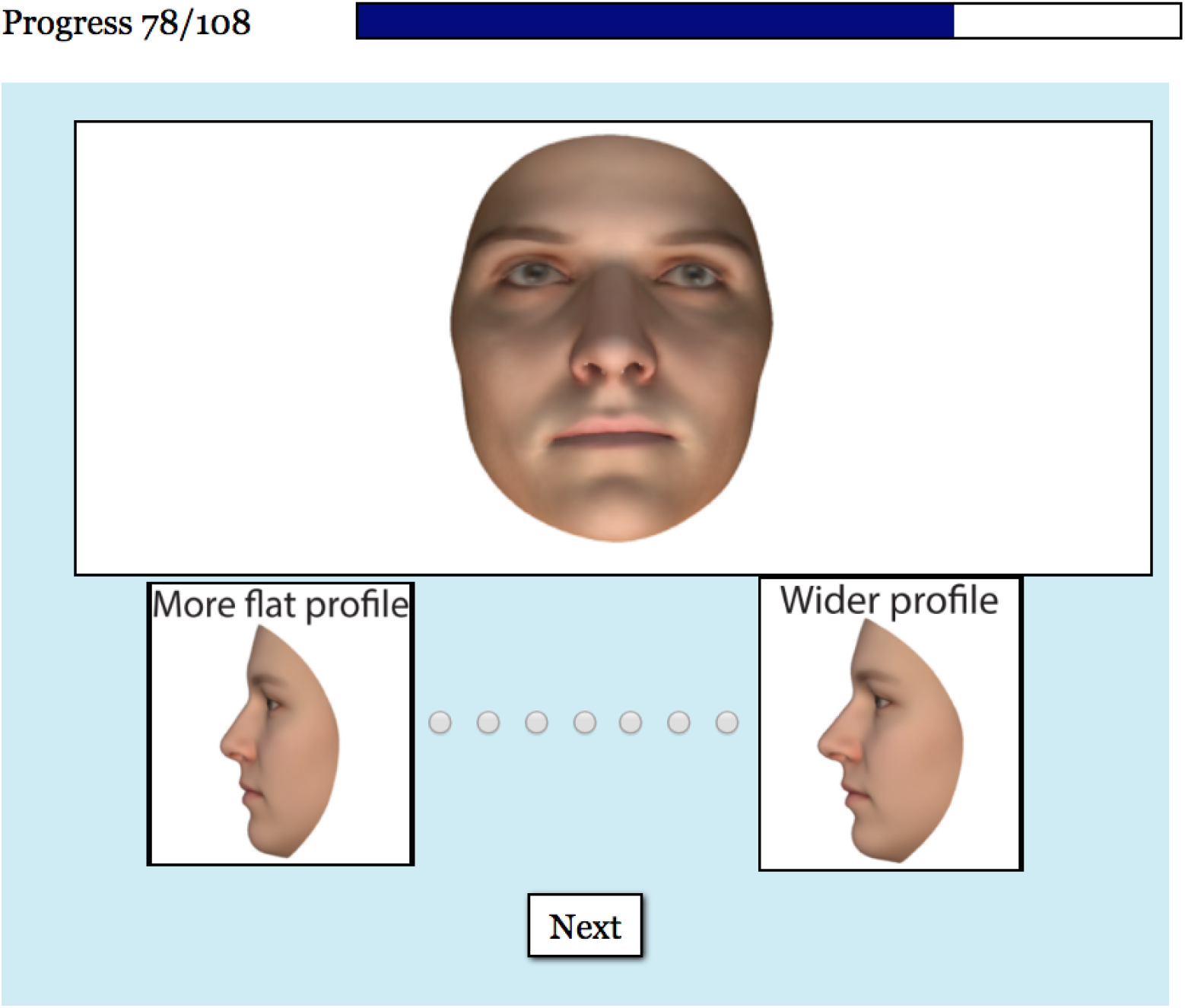
Snapshot of a trial from the depth judgment experiment.

**Extended Data Fig. 13:**
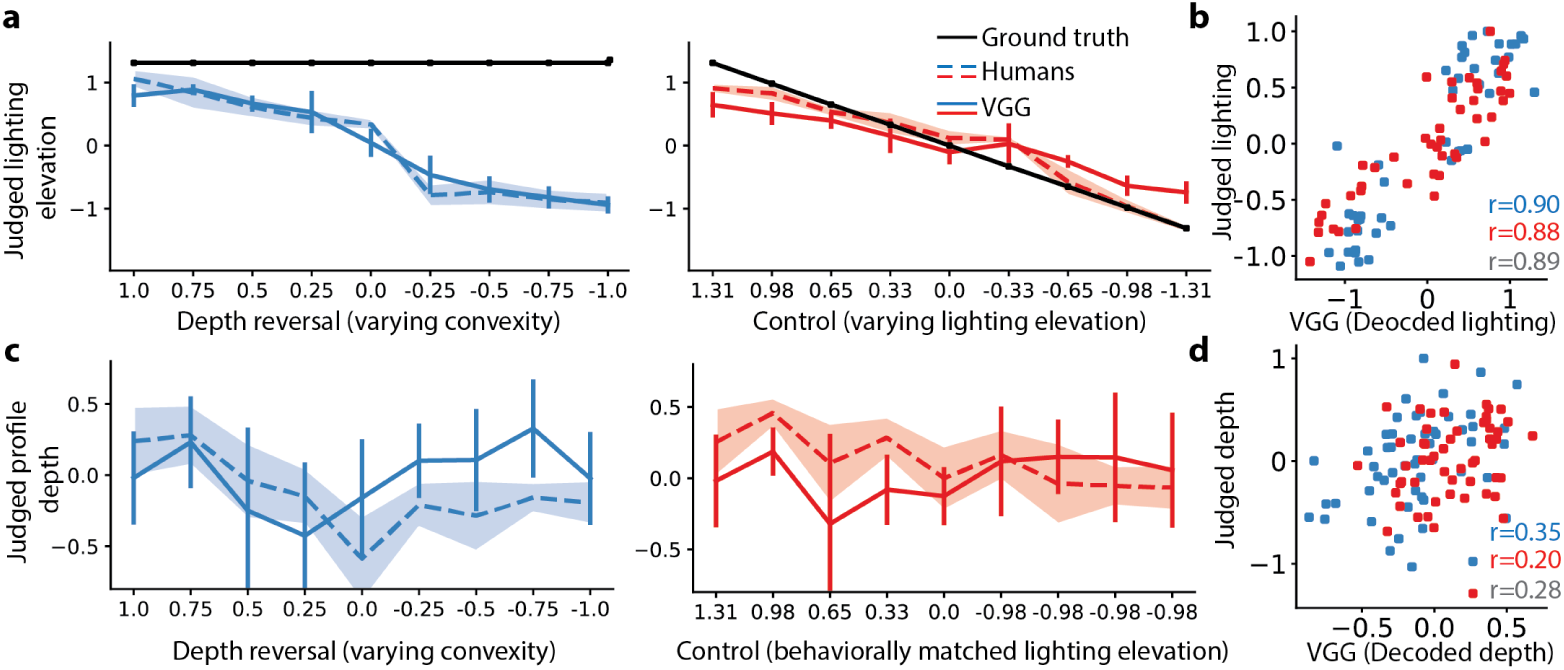
Decoding lighting elevation and profile depth from the VGG network. **a**, Normalized average light source elevation judgments of the depth-suppression group (left), the control group (right), lighting elevation values decoded from VGG, and the ground truth light source location. **b**, Average human judgments vs. lighting elevation decoded from VGG across all 90 trials without pooling to 9 bins. Pearson’s r values are shown for all trials (grey), control trials (red), and depth suppression trials (blue). **c**, Normalized average profile depth judgments of the depth-suppression group (left), control group (right) and profile depth values decoded from VGG. **d**, Average human judgments vs. profile depth values decoded from VGG across all 108 trials without pooling to 9 bins. Pearson’s r values are shown as in (**b**).

## Supplementary Information

### 1 Evaluation of alternative architectures and loss functions using the FIV-S image set

In this section, we summarize evaluations of various alternative network architecture and loss functions using the FIV-S image set (see Extended Data Fig. 4a). On the FIV-S images, EIG^−^ (Extended Data Figs. 4g-k) fit just as well as the full EIG model (Extended Data Figs. 4b-f), in both qualitative and quantitative terms. This confirms our expectation that the face segmentation stage is needed only to handle background clutter in the image, or hair or clothing that might occlude or distract from face shape and appearance, but that could also provide spurious cues to a familiar person’s identity.

When tested on the FIV-S, VGG-Raw showed less view-invariance overall (Extended Data Figs. 4r, s; *p <* 0.05 in comparison to VGG, Extended Data Figs. 4m, n). This relative lack of generalization across viewpoints suggests a form of dataset bias. On the other hand, VGG gave rise to patterns of results similar to its performance on the more natural FIV image set, with strong view-invariance in the top fully connected layers (Extended Data Figs. 4l-p). VGG’s top two layers remained similar to each other (Extended Data Fig. 4m), unlike both the neural responses (Fig. 2C, main text) and the EIG network (Extended Data Figs. 4c, h). Crucially, the training data used to finetune VGG matches EIG’s training data, which suggests that the superior fit of EIG to the neural data on the natural FIV faces, relative to VGG, is more likely a consequence of their respective targets as opposed to differences in training or test set distributions.

Two other VGG variants that further interpolated towards the EIG^−^ model were also tested to rule out possible alternative explanations for its lower fit, due to differences in architecture (Extended Data Figs. 4v-z; ID network, see below for its details), training loss functions (Extended Data Figs. 4aa-ae; Regress-ID,see below for its details), or alternative efficient analysis-by-synthesis methods with a learned image decoder based on a reconstruction loss (*β*-VAE and VAE-QN, see below for their details; Extended Data Fig. 5). Taken together, these results rule out the training data, the architectural differences between VGG and Alexnet, and the loss function as potential confounders. They positively support the inverse-graphics hypothesis for how the multistage inference network of primate face perception is organized: classic neural selectivity patterns across all three levels of ML/MF, AL and AM arise uniquely when a inference network is trained with targets that are 3D scene properties – that is, when the network is trained to infer the inputs to a causal generative model of observed face images.

#### ID network

This network allows us to rule out differences between the Alexnet and VGG architectures as a possible confound. Specifically, the architecture of the ID network was based on AlexNet similar to EIG except for its top layer (i.e., its TFCL), which was a 500-dimensional classification layer. We provide the details of this network’s architecture in Extended Data Table 3. This network was trained using the same generative model based dataset as VGG, minimizing a cross-entropy loss, but starting from randomly assigned initial weights.

#### Regress-ID network

Although the ID network matches EIG in training data and architecture, the loss function it optimizes is different from EIG. Moreover, ID network starts with random initial weights, unlike the EIG network. We built Regress-ID to equate the training loss function (MSE loss), the use of pretrained weights, and architecture to that of EIG. Moreover Regress-ID’s training set was the identical set of 200, 000 images we used for training EIG.

The Regress-ID network’s architecture is identical to the ID network except for it doesn’t have a classification layer (i.e., removing the TFCL layer in Extended Data Table 3 from the ID network gives the Regress-ID). We paired each image in the training set with an identity embedding vector representation as its target. The identity embeddings were obtained using the VGG network trained on identities from the generative model: For each image in the dataset, we recorded the SFCL activations of the VGG network. We trained the Regress-ID network to map images to these embeddings using stochastic gradient descent and a MSE loss.

#### VAE variants: *β*-VAE and VAE-QN

Finally, we built two VAE variants in order to test whether the exact form of the generative model mattered. EIG inverts a structured causal generative model based on a graphics program, however with VAEs generic function approximators (in this case convolutional neural networks) can be used to assimilate aspects of the generative model through learning. These models consist of an encoder (akin to an inference network) and a decoder (akin to a generative model). We equated these encoders to EIG’s in terms of their architectures and use of pretrained weights, similar to what we did for the Regress-ID network. (We also experimented with a pretrained VGG-Raw as encoder as well as with randomly initialized Alexnet architecture, neither of which led to any significantly different results to report here.) We tested *β*-VAEs^1^ and a modified Generative Query Network^2^, which are the specific variants of VAEs engineered to encourage learning disentangled representations from their inputs. In our case, this means disentangling scene extrinsic (e.g., pose) and scene intrinsic (shape and texture) elements of the scene.

To emphasize again, differently from EIG, these models don’t use a probabilistic graphics program as their generative model but instead learn one as part of training in the form of a convolutional neural network by minimizing a reconstruction loss, a loss function that is different from that of both EIG and VGG. (We provide its details below.) Overall, we found that these two models converge onto very different solutions from each other but neither captures the patterns in the neural data. One possible interpretation is that these models are not disentangling enough meaning that they are not capturing the overall causal structure underlying the images sufficiently. We now give the details of each the VAE variants.

*β-VAE.* The loss function that VAE optimizes, also referred to as the evidence lower bound (ELBO), is as the following^3^:

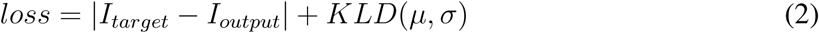

where *I*_*target*_ is the target image, *I*_*output*_ is the reconstructed image, *|I*_*target*_ *- I*_*output*_*|* is the reconstruction loss used in this study (i.e., l1 norm), and *KLD*(*μ, σ*) denotes a the Kullback-Leibler (KL) divergence with respect to the standard multivariate Gaussian distribution.

The reconstruction, *I*_*output*_ is obtained by inputting *z N* (*μ, σ*), a single sample from a multi-variate Gaussian distribution, to a decoder, where *μ* and *σ* are the outputs of an encoder that takes as input an image, *I*_*input*_.

As encoder, we used the Regress-ID architecture (including the pretrained AlexNet weights) except we replaced the SFCL layer with two fully connected layers each with 200 units. Following the re-parametrization trick in ^3^, these two layers represent parameters of a multivariate Gaussian distribution, *mu* and a transformation of *σ, log var*. *z* denotes a single sample from this distribution. The decoder (shared across *β*-VAE and VAE-QN) consisted of a series of 2D deconvolution and batch-normalization layers (Extended Data Table 4).

The *β*-VAE ^1^, the version of VAE we use, modifies the last term in Equation 1, *KLD*(*μ, σ*) to be *γ ∗* (*c - KLD*(*μ, σ*)), where *γ* and *c* are constants. The resulting equation is as the following.

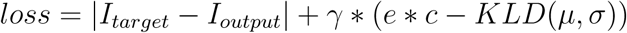

The hyperparameter *γ* controls the influence of the KLD associated loss. The hyperparameter *c*, referred to as controlled capacity, serves as an annealing parameter turning down the influence of KLD *e ∗ c* amount where *e* is the training epoch number.

In *β*-VAE training, *I*_*target*_ and *I*_*input*_ are identical images except *I*_*target*_ is downscaled to be 64×64 pixels, whereas *I*_*input*_ is 227×227 pixels. We trained *β*-VAE using the Pose dataset, which consisted of 2500 sets of images where each set contains 20 images of the same face identity with randomly sampled positions (total of 50000 images). Identities varied across sets. We use *γ* = 0.1, *C* = 100 and L1 distance as the reconstruction criteria between *I*_*target*_ and *I*_*output*_. We used Adam optimizer with a learning rate of 1 *∗* 10^−5^.

Despite our extended efforts – including experimenting with different encoders and decoders, trying a range of settings for *γ* and *C*, and using other datasets – this version of the VAE did not seem to learn to perform the task as indicated by the low view-invariance coefficients at layer SFCL, and more generally, this model did not give rise to patterns similar to those found in the neural data or in EIG.

##### VAE-QN

We also considered a second VAE variant, based on the recent Generative Query Network (GQN) that aimed to learn view-invariant scene representations as well as a neural graphics engine using a form of meta-learning ^2^. We formulated GQN for our purposes in the following way. Given an input image and a query pose, the network, like the *β*-VAE network, encodes the image and samples a *z* vector, but differently from *β*-VAE it also infers the pose of the face in the image. It then renders the scene captured in the *z* vector at the query pose. Following GQN, we reasoned that the network would need to learn to infer the intrinsic properties of an input face in order to be able to render it at random query angles. We refer to this model as VAE query network or VAE-QN in short.

The VAE-QN model used the same encoder architecture (and pretrained weights) as the *β*-VAE network for estimating *mu* and *sigma* and sampling *z*. It also used a parallel encoder for pose, which forks the output of the pre-TCL layer in the encoder (the convolutional layer just before TCL) and outputs a two dimensional pose vector, *P* = {*P*_*x*_, *P*_*z*_}, using a small convolutional network (architecture shown in Extended Data Table 5).

The basic decoder architecture followed that of The basic decoder architecture followed that of the *β*-VAE but required the following modifications to allow factoring in of the pose query. It included an initial fully connected layer that took as input a concatenated vector of *z* and *P* (vector of length 202) and outputed a 200 dimensional vector. This fully connected layer than was fed into the decoder architecture shown in Extended Data Table 4.

VAE-QN was trained following the same parameters as *β*-VAE, including *γ* = 0.1, *C* = 100 and L1 distance as a reconstruction criterion, and Adam optimizer with a learning rate of 1 *∗* 10^−5^.

We found that VAE-QN achieved a high-level of view-invariance by its SFCL, indicating that it learned the task. It also achieved a lower reconstruction loss on the validation set when compared to *β*-VAE. Indeed, we found reconstructions by this model to be less blurrier and more accurate than *β*-VAE. However, unlike EIG, this network gave rise to patterns that were similar to those found in the VGG network with high view-invariance and weaker mirror-symmetry in its FFCL.

### 2 Impact of pretrained weights and dropout rate

In this section, we report the impact that two key aspects of the training procedure have on the emergence of fully brain-like transformations: whether to start training from pretrained weights and the rate of dropout used during training. In each case, we evaluate how these aspects of training impact the EIG network in comparison to variants of the ID network by looking at the speed or efficiency of training and the resulting network similarity matrices.

#### Trade-off arising from the use of pretrained weights and training targets

The pretrained Alexnet weights provide the EIG network not only with a generic way to extract useful features from images ^4^ but it also gives a good initial representational state, in particular those of its TCL and FFCL, which contain the right qualitative structure (view-specificity in TCL and mirror-symmetry in FFCL) with respect to the neural data. Here we analyze the impact of starting training from this initial state, particularly how it interacts with the training targets implied by the classification vs. inference hypotheses.

On the classification hypothesis side, we include three variants of the ID network (ID-1, ID-2, and ID-3 networks) in addition to the VGG network. The ID network mentioned above (see also SI Section 1) shares the same architecture as EIG, but its weights are initialized randomly from scratch. Here, instead, the three variants of the ID network all start training from the pretrained weights, like the EIG network. All three networks were trained using the same generative model based dataset as the VGG network until an early stopping criteria on a shared validation set was triggered.

In the ID-1 network, we finetuned the layer SFCL starting from its pretrained weights and trained the layer TFCL starting from random initial weights. All other weights in the network were assigned and fixed to their pretrained weights. The ID-2 network, like the EIG network, updated the top four layers, including finetuning layers TCL, FFCL, and SFCL starting from their pretrained weights and training TFCL starting from random initial weights. The ID-3 network also updated all four layers like the ID-2 network but also included a high dropout rate (i.e., with probability 0.5) between each of the fully connected layers.

The ID-1 network’s FFCL – note that it is identical to that of the pretrained Alexnet by design – show a high level of mirror-symmetry (Extended Data Fig. 6a), consistent with the selectivity patterns seen in the neural data. However, this representational result comes at the cost of performing the task much worse. Its view-invariance coefficient is very low (Extended Data Fig. 6b) and its final average validation loss is very high in comparison all other models we considered. These indicate that the network cannot learn to perform the task of person identification much beyond the chance level. Consistent with this, its behavioral fits were poor (Extended Data Fig. 6c) including in Exp. 1 which does not require any particular qualitative generalization.

Overall, a strong pattern of trade-off emerged between performance and behavioral fits on one side and neural fits on the other: Across ID-1, ID-2, ID-3, and VGG networks, the better a network performed the classification task (Extended Data Fig. 6b) and the better it fitted the behavioral data (in particular Exp. 1; Extended Data Fig. 6c), the less it was mirror symmetric (Extended Data Fig. 6a). That is, through training, only by over-writing the existing mirror symmetry in FFCL of the pretrained weights the classification networks come to perform their corresponding task well. We illustrate this for ID-3 network in Extended Data Fig. 6d (right column) by plotting the mirror symmetry and view-invariance scores for its FFCL and SFCL layers throughout training.

This trade-off does not apply to the EIG network. Mirror-symmetry in EIG’s *f*_4_ layer increases with further training (Extended Data Fig. 6d, left column), and significantly exceeds the mirror symmetry found in the pretrained Alexnet weights (*p <* 0.001; EIG vs. ID-1 as shown in Extended Data Fig. 6a). Moreover, view-invariance converges to a high value in EIG’s *f*_5_ layer with further training (Extended Data Fig. 6b, and 6d left column). Finally, the EIG network gives overall better fits to the behavioral data (Extended Data Fig. 6c). In short, these results indicate that mirror-symmetry gets in the way of performance in the classification networks whereas it facilitates and is facilitated by training on the latent variables as targets in the EIG network.

#### Pretraining is not necessary, but it improves efficiency

Is pretraining necessary for a full set of brain-like transformations, including an intermediate mirror-symmetry stage, to emerge in the EIG network? To answer this question, we trained the EIG network starting from random initial weights as opposed to transferring weights from the pretrained Alexnet. We found that this training substantially reduced mirror-symmetry in the hidden full connectivity layer (*f*_4_, Extended Data Figs. 7a-d). However, when we trained a larger network with an additional hidden layer of full connectivity (*f*_4_*e* Extended Data Fig. 7e), we found that it recovered a significant level of mirror-symmetry at that extra hidden layer (Extended Data Figs. 7f, g). We also found that it took approximately five times longer to train these networks: three to five hundreds more training epochs were required to attain a loss value comparable to that of 75 epochs-long training with the original EIG network. Training of the network with two hidden full connectivity layers took about hundred more training epochs to reach the same levels of validation loss as the network with one hidden full connectivity layer. Even though longer training times are expected for networks starting from random initial weights, these results indicate that the original EIG training is efficient in two ways: (1) it reduces the number of hidden layers required for recapitulating the full set of of brain-like transformations, and (2) it trains much faster.

#### Number of hidden full-connectivity layers in EIG

We evaluated different architectures by varying the number of hidden full-connectivity layers before the scene properties layer to be zero, one (the standard EIG architecture; Extended Data Table 2), or two, meaning that the total number of full-connectivity layers could be one, two, or three. We found that with no hidden full-connectivity layers, training was not faster despite the smaller total number of parameters needed to be trained and the network did not give rise to an intermediate stage mirror-symmetry layer. Instead, it transformed directly from view-based representations in TCL to view-invariant representations in the FCL.

We found that with one hidden full-connectivity layer, a full set of brain-like transformations emerged provided that training started with the pretrained weights in Alexnet (e.g., Extended Data Fig. 4c; and see the subsection above). Finally, we found that with two hidden full-connectivity layers, an intermediate mirror-symmetry stage was always recovered independently of whether training started from scratch (Extended Data Fig. 7f) or from pretrained weights in Alexnet. In sum, for a full set of brain-like transformations to emerge, these results suggest that an Alexnet-like architecture is needed with multiple convolutional layers followed by at least one hidden full-connectivity layer before the scene properties layer.

#### Impact of dropout rate on the EIG network

The use of dropout in the ID networks (e.g., ID-3 vs ID-2 networks in Extended Data Fig. 6) improves performance while reducing mirror symmetry. Here we explore how the rate of dropout impacts the learned representations in the EIG network. Similar to the ID networks, we found that high dropout rates (e.g., dropout with probability 0.5, but not with probability 0.1 or 0.01) during any face-specific training (either finetuning or training from scratch) reduced mirror symmetry in the EIG network. However, unlike the ID networks, it did not lead to better behavioral fits nor any performance benefits in terms of the validation loss. Instead, using a high dropout rate resulted in a five-fold slow-down in the learning speed. (The number of epochs required to reach a certain level of validation loss were 373 epochs and 75 epochs for training with and without dropout.)

Note that during training, dropout randomizes the set of weights across forward calls breaking any potential symmetry of weights between pairs of mirror-symmetric poses. In contrast to EIG’s tolerance of low dropout rates, a perhaps more stringent requirement is observed in the theory developed by Leibo et al.^5^ where obtaining mirror-symmetry required squaring of the odd (but not even) PCA components, akin to using the same exact weights twice.

#### Training set with mirror-symmetric views

Finally, another requirement of the theory developed by Leibo et al.^5^ is a dataset containing mirror-symmetric views of the same identity. We found that this requirement was largely optional in EIG, with not including reflections of images over the vertical midline during training still resulted in mirror symmetry albeit at a somewhat reduced level.

### 3 Functionally interpreting ML/MF and *f*_3_ using the generative model

Albedos and normals for each of the 25 person identities in the FIV image set are approximated using EIG and the generative model. The 3D shape and texture properties for each frontal-pose FIV image are inferred using EIG (outputs at *f*_5_). Given the resulting 3D meshes, we obtained the face proper regions by masking out the neck, ears, and hair from the resulting 3D meshes for each identity. Using the generative model, we rendered the 2.5D components of each of the masked meshes at the 7 mean pose values underlying the extrinsic scene parameter distribution for the FIV-S image set (Extended Data Table 1). Finally, we adjusted the size and location of faces in the images using the same normalization procedure as the attended images.

We hypothesized that the random variables in the generative model (Fig. 1b main text and Extended Data Fig. 1a) that are hierarchically below the 3D scene properties each provide a conditional independence stage that could be exploited by ML/MF or AL. We tested this hypothesis using the similarities arising from each of the potential conditional independence stages (Fig. 3a, main text): raw input images, attended images, and 2.5D components including albedos and normals. The attended images and 2.5D components are both better accounts of ML/MF than the raw images (*p <* 0.001; raw images *r* = 0.29[0.24, 0.35], attended images *r* = 0.52[0.47, 0.57], albedos 0.60[0.56, 0.64], and normals 0.63[0.59, 0.67]) with the 2.5D components providing a significantly better account than the attended images (*p <* 0.001 for each 2.5D component). However, *f*_3_ continued to provide a better account of ML/MF than the 2.5D components (*p <* 0.001).

Similar to our ML/MF results (Fig. 3b, main text), we found that *f*_3_ similarity matrix itself was highly correlated with the 2.5D components to a much better degree than the raw input images (albedos 0.83[0.81, 0.85], normals 0.84[0.82, 0.86], raw images 0.36[0.28, 0.44]; *p <* 0.001 for each comparison), with the exception that attended images correlated with *f*_3_ as highly as the 2.5D components (0.86[0.85, 0.87]; Extended Data Fig. 8a). To better understand *f*_3_ and its relationship to attended images, we used the FIV-S-2 image set consisting of higher image-level variability which allowed us to tell apart attended images from the 2.5D components. Unlike the attended images, albedos and normals continued to correlate consistently well with *f*_3_ (raw images 0.25[0.23, 0.29], attended images 0.46[0.44, 0.50], albedos 0.85[0.82, 0.86], normals 0.87[0.85, 0.88]; Extended Data Fig. 8b). These results collectively suggest that both ML/MF and *f*_3_ can be understood as 2.5D-like surface representations and also suggests use of image sets with broader image-level variability in future experiments for better understanding ML/MF computations.

#### Linear decodability of the 2.5D-like representations

We consistently find that VGG and its variants trained to estimate face identity (Fig. 2j-m, Extended Data Fig. 2a-j, Extended Data Fig 4l-ae) do not produce an AL-like mirror symmetric representation distinct from both the 2.5D-like representation in ML/MF and EIG-*f*_3_ and the 3D scene property representation in AM and EIG-*f*_5_; instead, all fully connected layers of these networks have similar responses with strong viewpoint-invariant identity coding from the first fully connected layer (FFCL) upwards. To explain this, we hypothesized that these discriminatively trained networks are performing a fundamentally different computation in their hidden layers than EIG and the face-patch circuitry: While EIG and the ventral stream appear to need a distinct hidden-layer transformation to solve the nonlinear mapping from 2.5D surface components to 3D object properties – our interpretation for the function of AL and EIG-*f*_4_ – the identity classification task that VGG and its variants are trained for might be linearly solvable from the high-level image features computed in these networks’ top convolutional layer (TCL), with no need for further nonlinear transformations.

To test this hypothesis, we attempted to linearly decode identity on the FIV faces from each of the models: layers TCL, FFCL, and SFCL in VGG and layers *f*_3_, *f*_4_, *f*_5_ in EIG. Specifically, we trained a one-layer linear-softmax classification network on ML/MF, and on the max-pooling outputs of EIG-*f*_3_ and VGG-TCL, to decode all 25 FIV identities. We split the 175 FIV images to 6 poses (6 *×* 25 = 150 images) for training and 1 pose (25 images) for testing with averaging results across all 7 possible splits. Extended Data Fig. 9a shows the held-out test performance of the linear classifier. All layers in both networks gave rise to above chance (4%) decoding performance, but we found far better decodability of identity in the VGG network, with its TCL representation already achieving near FFCL and SFCL performance. In contrast, in the EIG network identity wasn’t nearly as linearly decodable initially at its *f*_3_ layer but increased to a comparable level of performance as the VGG network by layer *f*_5_ (Extended Data Fig. 9a left panel). These results support our conjecture that the face identity might be linearly solvable from the TCL representations of the VGG network and its variants, without a need for further nonlinear transformations.

We also tested whether a nonlinear transformation on top of the 2.5D-like representations –e.g., the layer *f*_4_ in the EIG network– are required for mapping these representations to 3D object properties. We attempted to linearly decode the shape and texture properties of the FIV images –approximated using the EIG network as its layer *f*_5_ outputs given the 175 FIV images– based on both models, layers TCL, FFCL, and SFCL in VGG and layers *f*_3_, *f*_4_, and *f*_5_ in EIG. We performed linear regression using the partial least squares (PLS) method with 33 retained components ^6, 7^. We split the 175 FIV images to 6 poses (6 *×* 25 = 150 images) for training and 1 pose (25 images) for testing with averaging results across all 7 possible splits. Extended Data Fig. 9b shows the goodness-of-fit *R*^2^ values on the held-out test sets. We found that these shape and texture vectors were not linearly decodable from any of the VGG layers, whereas it became increasingly more decodable in the EIG network from layer *f*_3_ to *f*_4_. Notably, the intrinsic scene properties (i.e., the shape and texture properties) were much less linearly decodable at layer *f*_3_ when compared to layer *f*_5_ indicating that indeed the transformation from 2.5D-like representations to 3D scenes requires some nonlinear transformation.

### 4 Neural data

The neural experiments and the data presented in the main text were originally reported in ^8^.

#### Stimulus and experimental procedure

The neural experiments used the FIV image set. FIV included images of 25 person identities with each identity viewed at 7 different head orientations: left-profile, left-half-profile, straight, right-half-profile, right-profile, upwards, downwards. (The original recordings also used an 8th viewing condition,the back of the head, but we didn’t analyze the corresponding data in this study).

Images were shown in a rapid serial presentation mode with 200 msec on-time followed by 200 msec blank screen with gray background. Images were presented centrally and subtended an angle of 7°. Monkeys were given a juice reward for maintaining fixation at the center of the screen for 3 seconds.

#### Neural recordings

Single-unit recordings were made from three male rhesus macaque monkeys (*Macaca mulatta*). Before the recordings, face-selective regions in each subject were localized using functional magnetic resonance imaging (fMRI). The face-selective regions were determined as the regions that were activated more to faces in comparison to bodies, objects, fruits, hands, and scrambled patterns. Single-unit recordings were performed at four of the fMRI-identified face-selective patches, all in the inferior temporal cortex: middle lateral and middle fundus areas ML/MF, anterior lateral area, AL, and anteriormedial area, AM. Following the original study, we combined the responses from the regions ML and MF in our analysis due to their general similarity (referred to as ML/MF).

A single neuron was targeted at each recording session, in which each image was presented 1 to 10 times in a random order. Following ^8^, we only analyze responses of the well isolated units.

### 5 Supplementary psychophysics methods and model-free analysis

#### Experiment 1

A total of 48 participants were recruited over Amazon’s crowdsourcing platform, Mechanical Turk (one additional participant was eliminated due to performing at or worse than the chance performance, 50%). The task took about 10 minutes to complete. Each participant was paid $1.50 ($9.00*/*hour). All participants provided their informed consent and were at the age of 18 or older according to their self-report.

The average performance of participants was 66% with a standard deviation of 7%, a min value of 53%, and a max value of 78%. Two tailed t-tests revealed that the performance of the Experiment 1 participants were not statistically distinguishable from that of the Experiment 2 participants (*p* = 0.23) but both Experiment 1 and 2 participants performed more accurately than the Experiment 3 participants (*p <* 0.001 and *p <* 0.05). See the corresponding subsections below for further details of the accuracy distributions of Experiment 2 and 3 participants.

The average reaction time of participants was 1479 msecs, with a standard deviation of 626 msecs, a min value of 426 msecs, and a max value of 3754 msecs. Two-tailed t-tests revealed that the reaction time distributions were not statistically distinguishable from each other across all pairs of the three experiments (*p >* 0.2 in all pair-wise comparisons). See the corresponding subsections for the reaction time statistics of Experiments 2 and 3.

To examine effects of learning in each experiment, we considered a moving-window performance of the participants as the trials progressed from the 1st trial to the 96th trial (Extended Data Fig. 10). We used a window size of 10 trials and stride of 1. For each participant and moving-window index, we found that participant’s average performance in the next 10 trials including the current trial. Even though the data suggested some learning in the early trials in Experiment 1 (e.g., significant difference between the 1st and 10th windows’ performance, *p <* 0.05), there was no indication of learning in Experiments 2 and 3 (e.g., no significant difference between the 1st and 10th windows, *p >* 0.35 in both experiments).

#### Experiment 2

A total of 48 participants were recruited over Amazon’s crowdsourcing platform, Mechanical Turk (seven additional participants were eliminated due to performing at or worse than the chance performance, 50% and two other participants were eliminated because their average reaction times were very short, 19 msecs and 30 msecs, much less than even the perception-to-action cycle of the expert video game players, 100 msecs). The task took about 10 minutes to complete. Each participant was paid $1.50 ($9.00*/*hour). All participants provided their informed consent and were at the age of 18 or older according to their self-report.

Participants’ average accuracy was 64% with a standard deviation of 7%, a min value of 52%, and a max value of 80%. Their average reaction time was 1542 msecs, with a standard deviation of 564 msecs, a min value of 151 msecs, and a max value of 3716 msecs.

#### Experiment 3

A total of 44 participants were recruited over Amazon’s crowdsourcing platform, Mechanical Turk (12 additional participants were eliminated due to performing at or worse than the chance performance, 50%, and one other participant was eliminated because their average reaction time was very short, 16 msecs, much less than even the perception-to-action cycle of the expert video game players, 100 msecs). The task took about 10 minutes to complete. Each participant was paid $1.50 ($9.00*/*hour). All participants provided their informed consent and were at the age of 18 or older according to their self-report.

Participants’ average accuracy was 61% with a standard deviation of 6%, a min value of 51%, and a max value of 73%. Their average reaction time was 1403 msecs, with a standard deviation of 472 msecs, a min value of 508 msecs, and a max value of 2792 msecs.

#### Bootstrap procedure

In order to quantify the correlations between the models’ predictions and the data, we sampled whole subject responses with replacement. We generated 10, 000 such boot-strap samples. All p-values were estimated using direct bootstrap hypothesis testing ^9^.

#### Lighting elevation judgment task

A total of 60 participants were recruited over Amazons crowd-sourcing platform, Mechanical Turk. The task took about 10 minutes to complete. Each participant was paid $1.50 ($9.00*/*hour). Half of these participants participated in the light source elevation condition, and the other half participated in the depth-suppression condition. The experimental procedure was identical between the two groups.

Before the beginning of the experimental trials, both groups of participants were instructed that they would see images of faces that could be lit anywhere from the top of the face to the bottom of the face using an illustration of the range of possible scene lighting conditions (Extended Data Fig. 11a). An example trial from the lighting source elevation condition is shown in Extended Data Fig. 11b.

#### Face depth judgment task

A total of 40 participants were recruited over Amazons crowdsourcing platform, Mechanical Turk. The task took about 15 minutes to complete. Each participant was paid $2.00 ($8.00*/*hour).

#### Linearly decoding lighting elevation and face depth from the VGG network layers

The VGG network doesn’t explicitly infer the lighting elevation or the profile depth of a face given an image. But it is possible that the representations it learns in its intermediate stages can support linearly decoding these latent variables. To test this possibility and therefore to evaluate the classification hypothesis on the hollow-face effect, we attempted to linearly decode lighting elevation and profile depth of a face from each of the VGG network layers.

We attempted to learn a linear mapping (using a fully-connected layer) from each of the three layers of the VGG network (TCL, FFCL, and SFCL) to lighting elevation based on an image set of 10000 face image-lighting elevation (*L*_*z*_) pairs drawn from the generative model. For both target variables (lighting elevation and profile depth which is described below), we trained the newly added linear layer using stochastic gradient descent while freezing weights in the rest of the network. We report results based on the layer where lighting elevation was most accurately decodeable (FFCL; decodings based on this layer also correlated best with the data). We found that the lighting elevation was linearly decodeable (Extended Data Fig. 13a control stimuli) and overall the resulting decoder matched human judgments to a good extent (Extended Data Fig. 13a, b), but it correlated (*r* = 0.89) with the data significantly worse when compared to EIG (*r* = 0.95, *p <* 0.01 two-tailed test for difference of two correlation coefficients). More important than this small difference in correlations with human judgments is the cause of the difference. In the illusory condition, both EIG and humans show a sharp nonlinear change in judged lighting direction between flat faces and the first step of concave faces (Fig. 5a left panel, main text). VGG, in contrast, shows a much more gradual, almost linear trend at the same point, and has much greater variance (larger error bars) in its predictions at this step and adjacent steps (Extended Data Fig. 13a left panel). This suggests that VGG has not captured the interaction between lighting and shape perception that we see in humans and EIG, and which appears to be characteristic of the approximate inverse graphics solution for faces, but perhaps is picking up on some weaker, less specific statistical correlate of lighting direction that can be decoded in images more generally.

We also attempted to learn a linear mapping from each of the same three layers in the VGG network to profile depths of faces using a segmented version of the same 10000 images as inputs and profile depths as outputs. For each face, we calculated its profile depth the same we did it for the EIG network (and described above): use the generative model to first obtain its 3D shape and assign a depth value as the average displacement of the three key points in the face – nose, left cheek, and right cheek– with respect to the mean face. We report results based on the layer where profile depth was most accurately decodeable (TCL). We found that the decoded profile depths did not capture the trends in the data (Extended Data Fig. 13b; overall correlation *r* = 0.28) as well as EIG did (*r* = 0.60, *p <* 0.001 two-tailed test for difference of two correlation coefficients). We also found that overall correlation with judged profile depths was slightly (but not significantly) higher with profile depths decoded from another layer of the network (SFCL; *r* = 0.35, *p* = 0.63 two-tailed test for difference of two correlation coefficients).

